# The role of intrinsically disordered domains in regulating G protein coupled receptor signaling

**DOI:** 10.1101/2025.09.10.675278

**Authors:** Jun Xu, Ruoyi Qiu, Alexander M. Garces, Harald Hübner, Xinyu Xu, Dorothee Weikert, Peter Gmeiner, Michael T. Lerch, Brian K. Kobilka

## Abstract

The α_2A_ adrenergic receptor (α_2A_AR) is a clinically important target for various diseases including hypertension, diabetes and chronic pain. Here, using single-molecule fluorescence resonance energy transfer imaging, we show how agonist-specific activation dynamics in both structured transmembrane domain (TMD) and intrinsically disorders regions (IDRs) of α_2A_AR lead to diverse signaling profiles. Through seven pairs of strategically designed fluorophore labels, we systematically investigate the real-time conformational changes of α_2A_AR. Our study reveals unique TM6 dynamics in α_2A_AR, featured by a high energy barrier for agonist-induced outward movements essential for activation. In contrast, we identify agonist-specific conformational dynamics of a partially disordered extracellular loop (ECL2), highlighting its role as a dynamic regulatory module that controls receptor function. Moreover, we characterize the conformational landscapes of the long third intracellular loop (ICL3), revealing its compact structural features and membrane-proximal localization in the basal state, where it acts as a negative allosteric regulator in transducer coupling. Furthermore, we identify multiful functional sub-states of ICL3 that are dynamically modulated by both kinase phosphorylation and drug efficacy. These findings offer previously underappreciated structural and dynamic insights into α_2A_AR function governed by both TMD and IDRs, and may open up new avenues for the development of better therapeutics.

---

G protein-coupled receptors (GPCRs) regulate a wide range of cellular responses to various extracellular stimuli, making them a major target class for drug development for a multitude of human diseases ^1^. Advances in structural biology have offered valuable molecular insights into GPCR mechanisms and functions by capturing high-resolution structures of many receptors in complex with various ligands and signaling proteins ^2^. The advent of AlphaFold has further transformed the field by offering remarkably accurate predictive models for nearly all GPCRs ^3^. However, to attain a complete understanding of the functional complexity of GPCRs, including G protein coupling specificity, biased signaling and allosteric regulation, it’s essential to complement static structural snapshots with insights into the dynamic properties of these receptors and their transducer complexes ^4^. Moreover, GPCRs feature multiple intrinsically disordered regions (IDRs), such as the N-terminus, the carboxyl terminus and the third intracellular loop (ICL3). Given their intrinsic dynamics and flexibility, these regions are often truncated or there is a lack sufficient density to discern their conformations in structural studies employing X-ray crystallography or cryo-EM ^5^, and AlphaFold is not yet able to accurately predict the conformations of these dynamic regions. Accumulated evidence suggests that these IDRs play important roles in allosterically modulating GPCR function ^6^. However, the underlying molecular mechanism remains unclear due to the experimental challenges in investigating membrane protein IDRs.

The α_2A_ adrenergic receptor (α_2A_AR) is a prototypical class A GPCR that is distributed throughout the central nervous system (CNS) and peripheral tissues. It plays a pivotal role in regulating a number of physiological functions including neurotransmitter release, vascular tone regulation, insulin secretion, and nociception upon activation by endogenous neurotransmitter norepinephrine (Norepi) and the hormone epinephrine (Epi) ^7^. In contrast to β2 adrenergic receptor (β2AR), which primarily couples to the stimulate Gs protein to increase cellular cAMP, the α_2A_AR primarily couples to the inhibitory Gi/o proteins, reducing cellular cAMP production (**Fig. 1a**). The agonists of α_2A_AR have been in clinical use for decades, primarily in the treatment of hypertension. The α_2A_AR is also a validated non-opioid target for pain relief. However, the sedative effect of traditional α_2-_agonists like clonidine and dexmedetomidine (Dex) restricts their broad use as antihypertensive drugs or analgesics ^7–9^. Previous studies have shown that some partial agonists or G protein biased agonists that can selectively activate Gi/o proteins while avoiding arrestin recruitment can separate the sedative effect from analgesia or antihypertension ^10,11^. Therefore, understanding the molecular basis for α_2A_AR ligand efficacies and signaling bias is of great interest. Notably, α_2A_AR possesses a long ICL3 containing approximately 150 amino acids (**Fig. 1a, Extended data Fig. 1a**), accounting for nearly one-third of the entire protein sequence. Previous functional and biochemical studies have revealed that the ICL3 of α_2A_AR is a substrate for GPCR kinases (GRKs) and β-arrestin binding to regulate downstream signaling (**Fig. 1a**) ^12–14^. Studies have also identified key regions in the ICL3 that serve to direct agonist-promoted receptor desensitization and that mediate interactions with different G protein subtypes ^15,16^. Thus, a deeper understanding of the conformation and dynamics of the ICL3 is necessary to provide insight into the mechanism underlying functional selectivity.

**Fig. 1.**
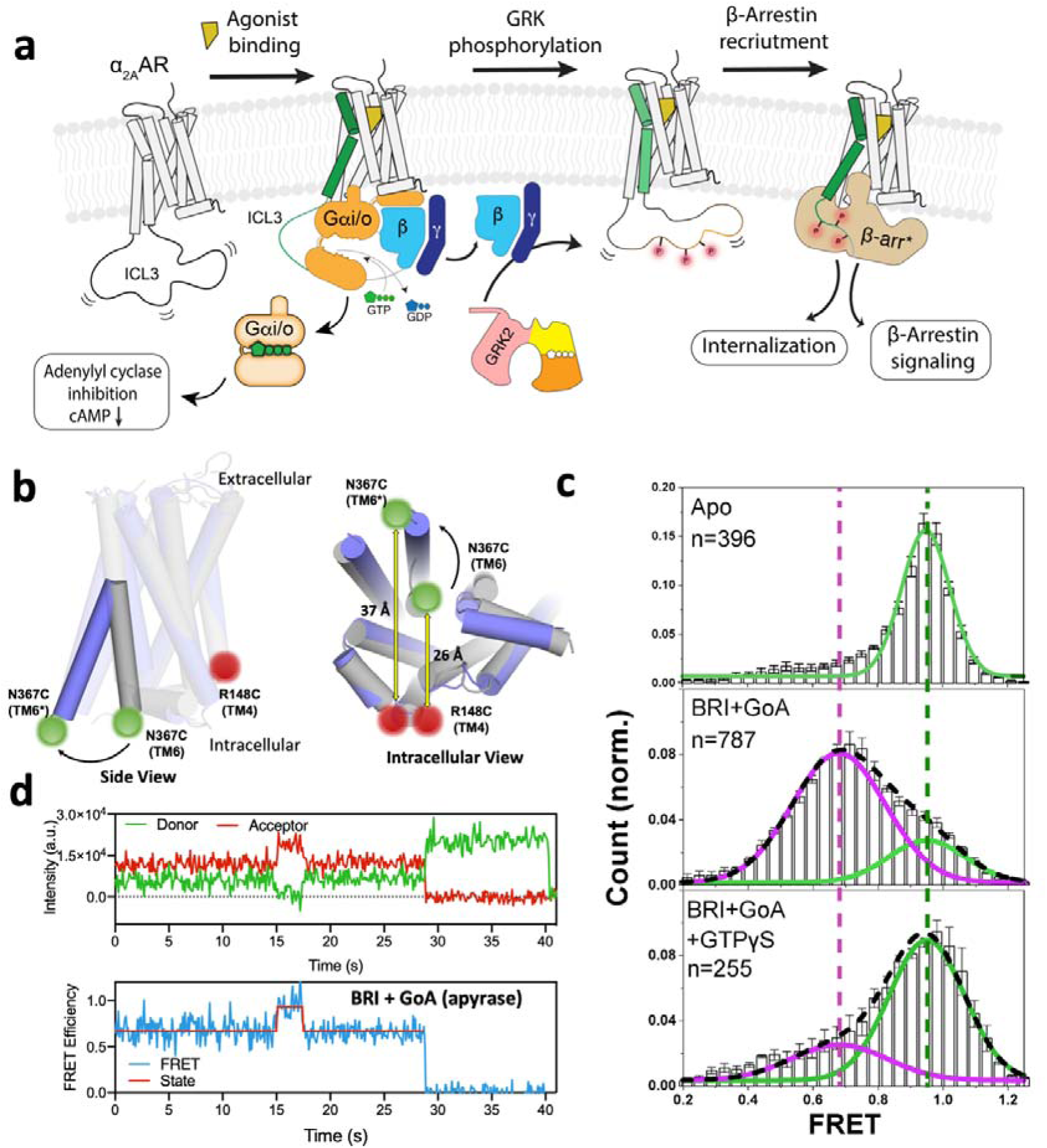
Modified receptor for smFRET investigation of α_2A_AR activation. (**a**) Schematic of α_2A_AR signaling pathways. (**b**) The outward movement of TM6 upon α_2A_AR activation. The inactive and active α_2A_AR were shown in gray and slate, respectively. (**c**) SmFRET distributions of TM4-TM6 sensor in the Apo, agonist (BRI) + GoA (apyrase) and BRI + GoA (GTPγS) bound states. Green and pink lines represent Gaussians fitted to the high-FRET inactive and low-FRET active states, respectively. Green and pink dashed lines indicate the distinct mean FRET values. Black dash line represents the cumulative fitted distributions. n represents the number of traces used to calculate the corresponding histograms. Data are mean ± s.e.m. from at least three groups of independent movies. Each group contains at least 3 movies. (**d**) Representative single-molecule fluorescence (donor, green; acceptor, red) and FRET time traces (blue) overlaid with predicted state sequence (red) showing transitions between inactive and active states. The trace was picked from agonist (BRI) + GoA^apyrase^ condition.

Here, using total internal reflection fluorescence (TIRF)-based single-molecule fluorescence resonance energy transfer (smFRET) imaging, we investigated the real-time conformational dynamics in different regions of α_2A_AR. A series of specifically labeled fluorophore pairs provide high-resolution information on conformational landscapes of TM6, the second extracellular loop (ECL2) and the unstructured ICL3 of α_2A_AR, and how the conformation and dynamics of these regions are regulated by five representative α_2A_AR agonists with distinct pharmacological profiles, by phosphorylation, as well as by transducer coupling. Combined with DEER spectroscopy, biochemical and cell signaling assays, these studies offer unprecedented molecular insights into the mechanisms by which the IDRs and transmembrane domain (TMD) concertedly govern drug efficacy and transducer specificity of GPCRs.

## Results

### Modified receptor for smFRET investigation of α_2A_AR activation

To site-specifically label the α_2A_AR with fluorescent probes through thio-selective chemistry, we first developed a minimal-cysteine α_2A_AR, in which three solvent-exposed cysteines were mutated to Ala or Ser (hereafter named α_2A_ARΔ3). The engineered α_2A_ARΔ3 exhibits wild-type functional properties (**Extended data Fig. 1b-c**). We next introduced two additional cysteines at the cytoplasmic ends of TM4 (R148C) and TM6 (N367C) to monitor the large outward movement of TM6, a well-known hallmark of GPCR activation (**Fig. 1b, Extended data Fig. 1a**). A similar labeling strategy has been proved to effectively report conformational dynamics for several other GPCRs including the β2AR, and μ-opioid receptor (μOR) ^17,18^. The construct (α_2A_ARΔ3-148/367, TM4-TM6 sensor) was purified and labeled with Alexa fluorophore 555(AF555) and 647 (AF647), the donor and acceptor fluorophores, respectively. The labeling specificity was verified by comparison with α_2A_ARΔ3, which showed negligible background fluorophore labeling under the same conditions (**Extended data Fig. 1d**).

For TIRF imaging, the labeled and N-terminal Flag-tagged receptor was immobilized to the surface of a polyethylene glycol (PEG) passivated coverslip through biotinylated anti-flag M1 fab, using a NeutrAvidin bridge (**Extended data Fig. 1e-g**). In the apo condition, the TM4-TM6 sensor exhibited a single Gaussian distribution with a high-FRET centered at ∼0.95 (**Fig. 1c**), which corresponds to the inactive TM6 conformation with close proximity to TM4. In an agonist (brimonidine, BRI) and GoA-bound complex treated with apyrase, we observed a predominant lower-FRET peak (∼0.67) (**Fig. 1c**), corresponding to the TM6 outward conformation upon G protein coupling. A small tail of high-FRET population likely results from receptors that did not couple with G protein and/or intrinsic dynamics within the nucleotide-free complex. Indeed, inspection of individual trajectories revealed transitions between the high- and low-FRET states which could be fitted with a two-state hidden Markov Model (HMM) (**Fig. 1d**). The histogram was therefore described as two Gaussian peaks (**Fig. 1c**). Addition of GTPγS to the α_2A_AR-GoA sample shifted the population predominantly back to the high-FRET state, suggesting the dissociation of the complex (**Fig. 1c**). Together, these data demonstrated the reliability of our smFRET design to examine the conformational changes of α_2A_AR during the activation process.

### Agonist-binding fails to stabilize TM6 outward movement

With the validated TM4-TM6 FRET sensor, we first examined the effects of different ligand efficacies on TM6 conformational dynamics. We selected five agonists, including the endogenous catecholamine agonist Norepi, two traditional imidazoline drugs dexmeditomidine (Dex) and moxonidine (Moxo), as well as two recently identified small molecule agonists 0172 and PS75 (**Fig. 2a**)^11^. We performed cell-based BRET assays to show that cysteine engineering has minimal effect on the functional profiles for all chosen ligands (**Fig. 2b, Extended data Fig. 1h-i**). As shown in Fig. 2b, Norepi is a full agonist in both G protein activation and arrestin recruitment; Dex and Moxo are full agonists in G protein activation while exhibiting partial agonism in arrestin recruitment; PS75 and 0172 are two G protein biased partial agonists with very low efficacy in arrestin recruitment. We first imaged the TM4-TM6 sensor in the presence of each ligand at saturating concentrations. However, we did not observe a significant change in the FRET distribution as compared to the apo condition, even for the high-efficacy agonist Norepi (**Fig. 2c**). Inspection of individual trajectories revealed relatively stable fluorescence and FRET under these conditions, and there is no significant shift of FRET peak centers (**Extended data Fig. 2a-b**). We initially suspected that this might be due to the large R0 (51 Å) for AF555 and AF647, such that the relatively small distance change induced by ligand alone could not be resolved due to the small FRET window (**Extended data Fig. 2c**). We then labeled the receptor with Cy3 and Cy7 which have smaller R0 (42 Å) and larger FRET window within the same distance range (**Extended data Fig. 2c**). Consistent with the smaller R0, we observed a lower-FRET peak centered at ∼0.6 for Cy3/Cy7 labeled receptors, but we still did not detect obvious change upon agonist binding (**Extended data Fig. 2d**). This is in contrast to what has been observed in previous spectroscopic studies of β_2_AR, where an agonist-dependent TM6 tilt to an active-like conformation was detected, even for partial agonists ^17,19^. To test whether this discrepancy originated because of either the different dye pairs or a different environment, we imaged the β_2_AR TM4-TM6 sensor which was purified in the same detergent environment and labeled with the same dye pair AF555/AF647. In line with published results ^17^, we observed a small but significant shift of the FRET center upon binding with the agonist isoprenaline (ISO), and Gs coupling further shifted the population to a fully active state at mid-FRET (**Extended data Fig. 2e-g**). These data indicate that the α_2A_AR has distinctive TM6 dynamic properties from what was observed in β_2_AR. Notably, a recent smFRET study on μOR also revealed that partial agonists have minimal effects on TM6 dynamics, with only highly efficacious full agonists inducing outward TM6 movement ^18^.

**Fig. 2.**
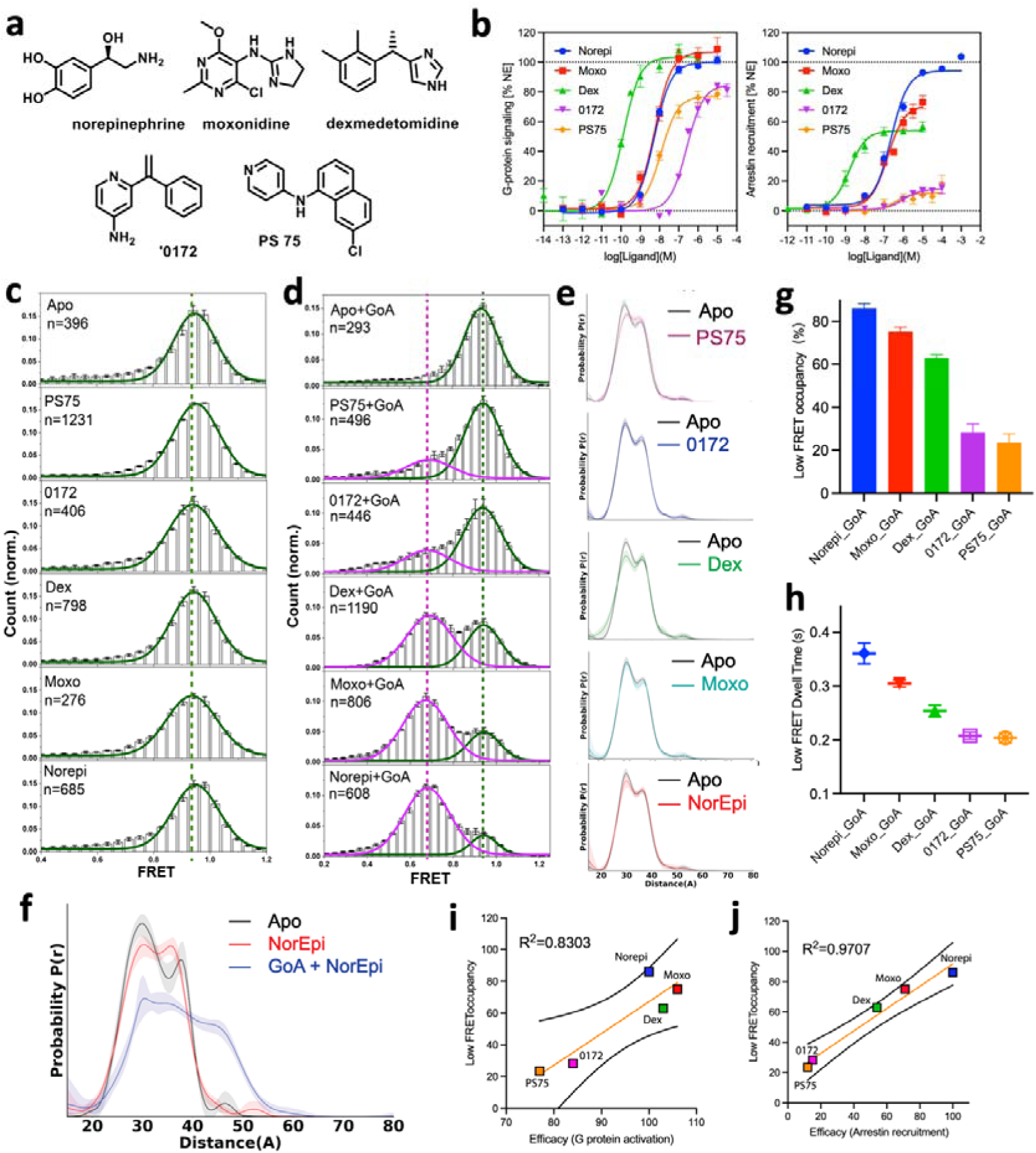
Conformational dynamics of TM6. **(a)** Chemical structures of selected ligands. **(b)** Signaling efficacy towards G protein activation and β-arrestin-2 recruitment for selected ligands. **(c-d)** SmFRET distributions of TM4-TM6 sensor bound to ligands only (**c**) or ligands + GoA treated with apyrase (**d**). Green and pink lines represent Gaussians fitted to the high-FRET inactive and low-FRET active states, respectively. Dashed green and pink lines indicate the distinct mean FRET values. n represents the number of traces used to calculate the corresponding histograms. Data are mean ± s.e.m. from at least three groups of independent movies. Each group contains at least 3 movies. **(e)** DEER distance distributions of the spin-labelled α_2A_AR TM4-TM6 sensor in different ligand conditions and (**f**) with GoA (Norepi+GoA^apyrase^) **(g-h)** Low-FRET occupancies (**g**) and dwell times (**h**) of TM4-TM6 sensor stabilized by different ligands and GoA. Data are mean ± s.e.m. from at least three group of independent movies. **(i-j)** The positive correlation of the low-FRET occupancy and ligand efficacy in G protein activation (**i**) and β-arrestin-2 recruitment (**j**).

To confirm the lack of an agonist-dependent shift in the conformational equilibrium of TM6 for the α_2A_AR, we performed double electron-electron resonance (DEER) spectroscopy experiments on the TM4-TM6 sensor spin-labeled with IA-proxyl. DEER provides a probability distribution of distances between spins and has high sensitivity in detecting conformational changes at Ångstrom resolution. The distance distributions of apo and agonist-bound α_2A_AR are very similar (**Fig. 2e, Extended data Fig. 3a**), in agreement with smFRET measurements. The distributions exhibit peaks at two distances (∼30 Å and ∼36 Å), which likely represent two distinct inactive conformations that lie in the same high-FRET (∼0.95) range (**Extended data Fig. 2c**). The active conformation (∼46 Å) was only observed in the presence of both agonist and GoA (**Fig. 2f, Extended data Fig. 3b**), which corresponds to the lower-FRET state (∼0.67). By contrast, agonist alone was found to induce a significant increase of active state population in previous DEER studies of β_2_AR ^19^.

Together, these results further suggest that, unlike β_2_AR, agonist alone is insufficient to stabilize outward motions of TM6 in α_2A_AR or the motions are too small to be detected by the techniques used here. This could be explained by different energy landscapes where the energy required for TM6 transition from an inward to outward conformation is higher in α_2A_AR than β_2_AR (**Extended data Fig. 3c**). Considering that α_2A_AR primarily couples to Gi/o while the β_2_AR selectively couples to Gs, we speculate that the distinctive conformational dynamics of TM6 may have a role in G protein coupling specificity. In line with this notion, cryo-EM structures revealed TM6 outward displacement in Gi/o coupled receptors is smaller than in Gs coupled receptors ^20–22^. Moreover, a recent smFRET study also showed quite distinctive TM6 dynamics between two chemokine receptors that couple different signaling proteins ^23^.

To further investigate ligand effects on conformational dynamics of TM6 in signaling complex, we imaged the TM4-TM6 sensor in the presence of different agonists and GoA. Although there wasn’t any detectable change in TM6 for agonist binding alone, GoA coupling shifted the population towards a low-FRET active state for all tested agonists at the expense of the high-FRET peak (**Fig. 2d, Extended data Fig. 3d**). No significant low-FRET population was observed when incubating GoA with Apo receptor, suggesting that agonist binding stabilized a pre-active state that facilitates GoA binding (**Fig. 2d**). Additionally, the occupancy and dwell times of the low-FRET active state varied depending on the ligand, following the order: Norepi > Moxo > Dex > 0172 and PS75 (**Fig. 2g-h, Extended data Fig. 3e**). This indicates distinct efficiency and dynamics of complex formation when bound to different agonists. Notably, the active state occupancy measured by smFRET was substantially lower than the G protein activation efficacy observed in cell-based BRET assays, particularly for 0172 and PS75, which exhibited ∼80% efficacies while only populating ∼25% in the low-FRET state (**Fig. 2b, g**). This suggests that even a small fraction of activated receptors can generate a significant cellular signaling response. The discrepancy might also reflect an intrinsic difference between the single-molecule and bulk assays. Nonetheless, there was an overall positive correlation between low-FRET occupancy and ligand efficacy in G protein activation (**Fig. 2i**). Of interest, we found that the GoA stabilized low-FRET occupancy also correlated well with ligand efficacies in arrestin recruitment (**Fig. 2j**), indicating a potential role of G protein activation in arrestin signaling pathway. Supporting this notion, previous functional studies revealed that α_2A_AR is predominantly phosphorylated by GRK2, and the phosphorylation requires the recruitment of GRK2 to the membrane via the dissociated Gβγ subunit of Gi/o ^24,25^. Thus, G protein activation may affect the subsequent arrestin recruitment to α_2A_AR through GRK2 phosphorylation.

### Ligand efficacy modulates conformational dynamics of ECL2

The agonist-specific complex formation efficiency and dynamics suggested that these ligands stabilized distinct ‘pre-active’ conformational states prior coupling to a G protein. To investigate the potential conformational differences upon binding with different agonists with smFRET, we developed a TM4-ECL2 sensor by introducing a cysteine mutation at Ala184 paired with the TM4 cysteine (α_2A_△3-148/184) (**Fig. 3a, Extended data Fig. 4a**). The ECL2 was not well resolved in both inactive crystal and active cryo-EM structures of α_2A_AR (**Extended data Fig. 4b-c**), suggesting high structural flexibility of this loop. Notably, the ECL2 of α_2A_AR exhibits an extended conformation in the Alphafold2 predicted model, in contrast to the folded conformation of the resolved part from determined structures, indicating potential large structural changes (**Extended data Fig. 4b-c**). In addition, ECL2 is closer to the orthosteric site and previous cryo-EM structures have revealed that the Ile205^45,52^ (ECL2) residue has direct interactions with most α_2A_AR agonists ^11,21^. Thus, the TM4-ECL2 sensor could be more sensitive to conformational changes associated with orthosteric ligand binding than the intracellular TM4-TM6 sensor.

**Fig. 3.**
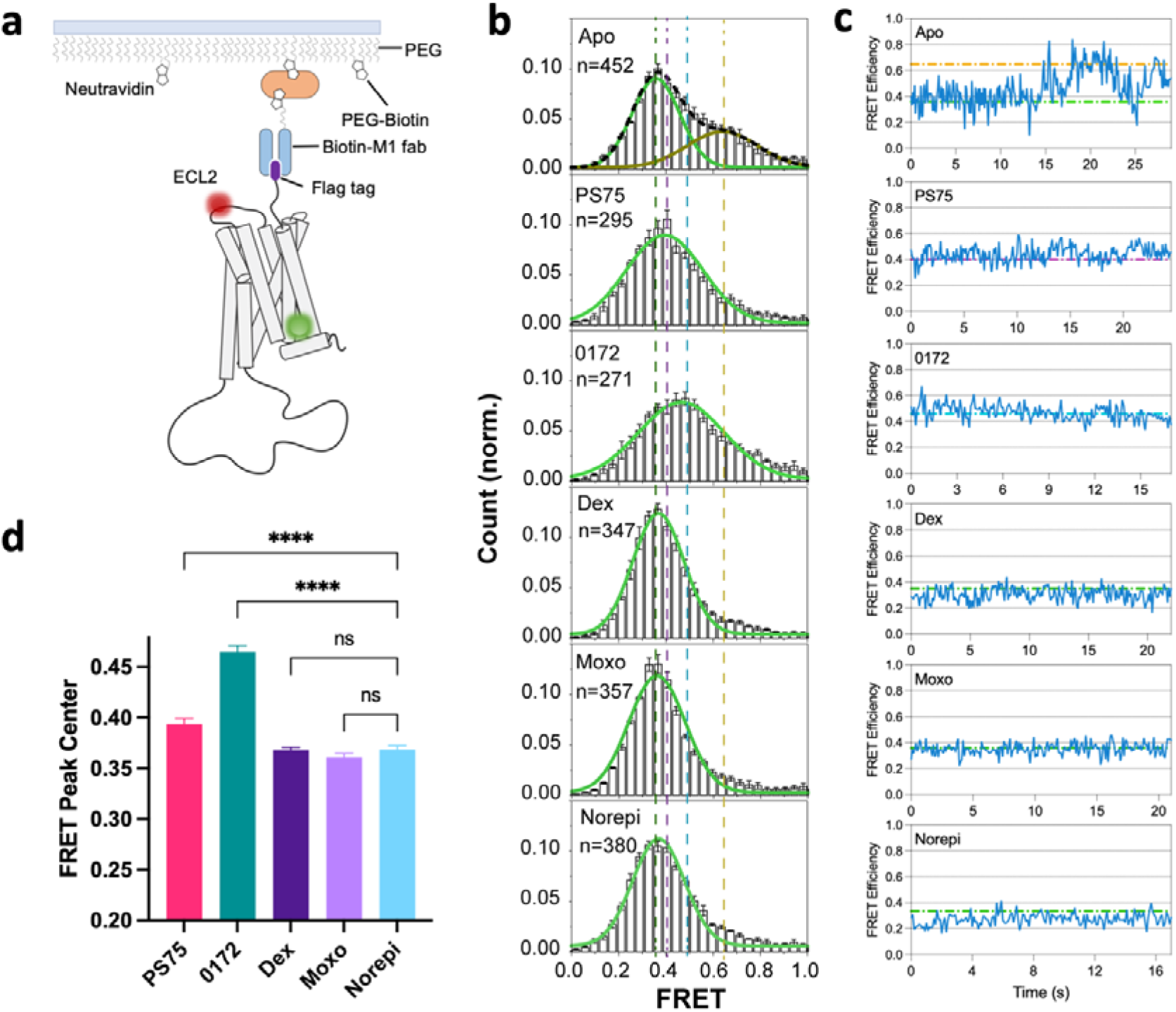
Conformational dynamics of ECL2. (**a**) Schematic of smFRET experiments for TM4-ECL2 sensor. (**b**) SmFRET distributions of TM4-ECL2 sensor in the presence of different ligands. Green and orange lines represent Gaussians fitted to the high-FRET inactive and low-FRET active states, respectively. Black dash line represents the cumulative fitted distributions. Colored vertical dashed lines indicate the distinct mean FRET values. n represents the number of traces used to calculate the corresponding histograms. Data are mean ± s.e.m. from at least three groups of independent movies. Each group contains at least 3 movies. (**c**) Example single-molecule FRET traces (blue) of TM4-ECL2 sensor are shown for each ligand condition. Colored dash lines indicate the distinct mean FRET values. The corresponding fluorescence traces for each condition were shown in Fig. S4D. Data are mean ± s.e.m. from at least three group of independent movies. (**d**) Comparison of FRET peak centers for TM4-ECL2 smFRET sensor in the presence of different agonists. Data are mean ± s.e.m. from at least three groups of independent movies.

Unlike the TM4-TM6 sensor, the TM4-ECL2 sensor exhibited a heterogeneous FRET distribution in the absence of ligand (apo), which populated two major states: a predominant low-FRET state (∼0.35) and a mid-FRET state (∼0.65), suggesting that the loop can adopt multiple conformations that move relative the TM domain with a slow time scale over 100 ms (**Fig. 3b**). We also observbed the transitions between low- and mid-FRET states from individual trajectories, highlighting the dynamic nature of ECL2 in the apo condition (**Fig. 3c, Extended data Fig. 4d**). Addition of the full agonists Norepi, Moxo and Dex resulted in a uniform FRET distribution centered at ∼0.35. Inspection of individual trajectories revealed relatively stable fluorescence and FRET, suggesting reduced conformational heterogeneity of ECL2 (**Fig. 3b-c, Extended data Fig. 4d**). By contrast, addition of the two weaker G protein biased agonists 0172 and PS75 led to relative broad FRET distributions centered between the low-FRET (∼0.35) and mid-FRET (∼0.65) (**Fig. 3b-d, Extended data Fig. 4d**). This is likely due to fast transitions between different states that could not be well resolved at current time resolution (100 ms), therefore resulting in a time-averaged intermediate FRET. It’s also possible that 0172 and PS75 stabilized different ECL2 conformations not observed in the apo and full agonists-bound conditions. Together, these results suggest ligands with different efficacies and bias can stabilize distinct conformational dynamics of ECL2. Notably, the coupling between ECL2 and signaling efficacy resembles what has been observed in other family A receptors such as the D2 dopamine receptor (D2R) and M2R, supporting the notion that ECLs play a role in modulating GPCR function and may provide a target for the development of functional selective drugs ^26–28^.

### Conformational dynamics of ICL3 in the apo state

In addition to extracellular loops (ECLs), intracellular loops (ICLs) also play crucial roles in modulating GPCR function. The ICL1 and ICL2 loops of GPCRs are relatively short and well-resolved in nearly all determined structures. Structures of GPCR-transducer complexes revealed direct involvement of ICL1 and ICL2 in transducer binding ^29^. In contrast, the ICL3 shows remarkable diversity in amino acid sequence and overall length, with only a few GPCR structures with short ICL3 being resolved ^6^. The long ICL3 of α_2A_AR was not resolved in all structural studies using full-length receptors due to their intrinsic flexibility and disordered nature ^11,21^. To better understand the structure-function relationship of ICL3, we characterized its conformational dynamics using smFRET. Given the extreme length of ICL3 in α_2A_AR, we introduced 5 cysteine mutations (E352C, R322C, D292C, G262C and T227C) evenly distributed throughout the ICL3 loop, starting from the bottom of TM6 to the intracellular end of TM5, each paired with the TM4 cysteine (148C) (**Fig. 4a-b**). We first imaged these ICL3 sensors in the absence of agonist to determine its conformational landscape in the apo state and found that different regions of the ICL3 displayed distinctive conformational states and dynamic properties (**Fig. 4c**).

**Fig. 4.**
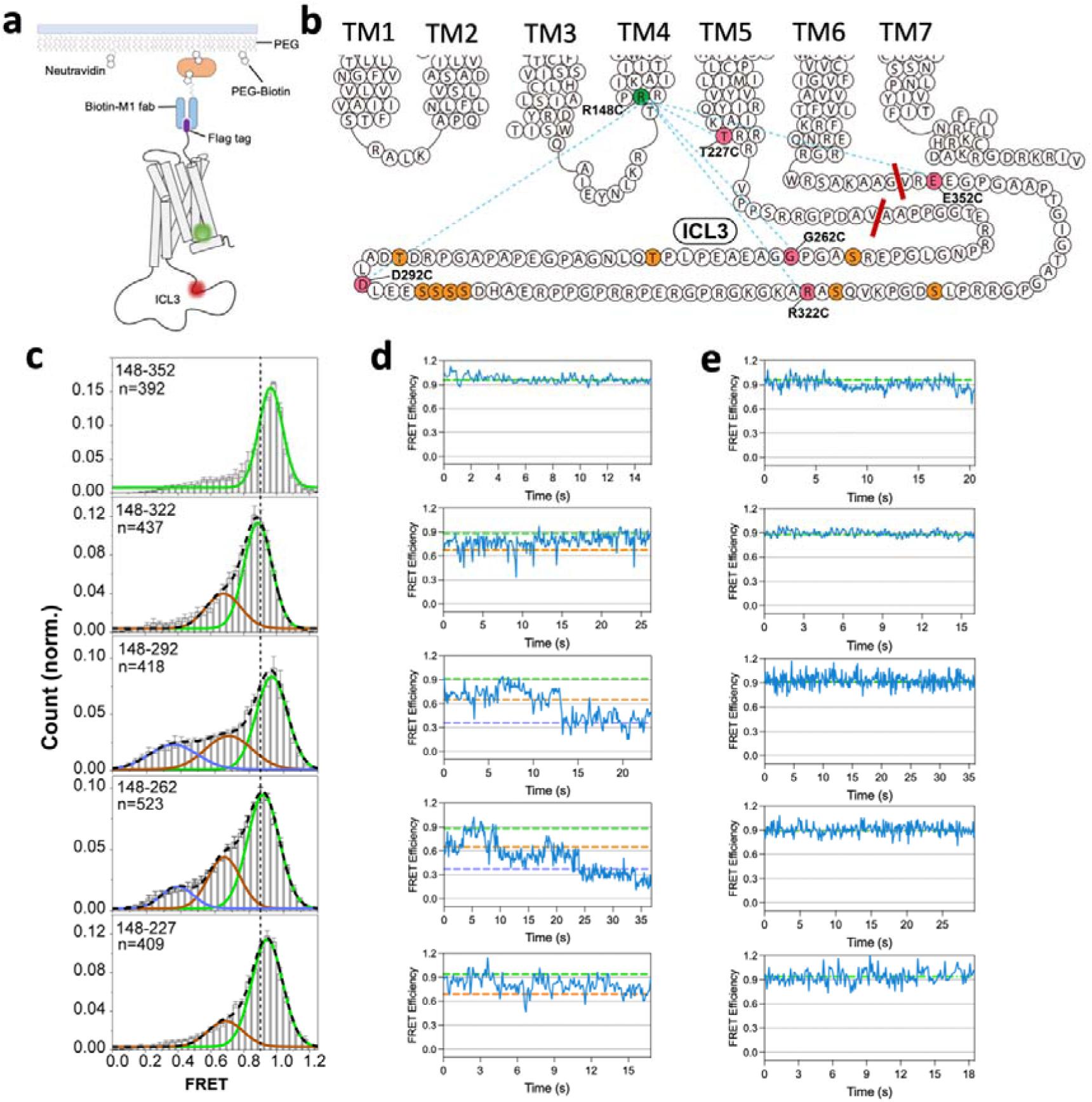
Conformational dynamics of ICL3 in the apo state. **(a)** Schematic of smFRET experiments for TM4-ICL3 sensor. **(b)** Positions of 5 labeling sites in ICL3 (red) and potential phosphorylation sites in ICL3 (orange). Red lines indicate the ICL3 truncation sites. **(c)** smFRET distributions of TM4-ICL3 sensors in the Apo state. Green, orange and slate lines represent Gaussians fitted to high-, mid- and low-FRET states, respectively. Black dashed line represents the cumulative fitted distributions. n represents the number of traces used to calculate the corresponding histograms. Data are mean ± s.e.m. from at least three groups of independent movies. Each group contains at least 3 movies. The vertical lines represent FRET 0.9. **(d-e)** Two example single-molecule FRET traces (blue) for in each condition are shown in (**d**). Colored dash lines indicate the distinct mean FRET values fitted by Gaussian.

When the FRET sensor is located at the C-terminus of ICL3 (148/352), which is ∼15 amino acids upstream of the intracellular end of TM6, we observed a single Gaussian distribution centered at FRET ∼0.95 (**Fig. 4c-e**), similar to that of the TM4-TM6 sensor. We started to see more heterogeneous conformational states at the 148C/322C sensor located further away from TM6 (∼45 amino acids upstream), as evidenced by the population at mid-FRET ∼0.65, in addition to the main population at high-FRET ∼0.95. We also observed clear fluctuations from high- to the mid-FRET state in individual trajectories (**Fig. 4d, Extended data Fig. 5a**). The FRET distribution became even broader for the 148/292 and 148/262 sensors, which are located in the middle region of ICL3 (**Fig. 4c**). In addition to the high-(0.9-0.95) and mid-FRET (∼0.65) populations, there appears an increased population at low-FRET state (∼0.4) for both sensors, indicating more complicated conformational dynamics. Inspection of individual trajectories also revealed conformational transitions among these three states (**Fig. 4d, Extended data Fig. 5a**). It is noteworthy that previous identified kinase phosphorylation sites (**Fig. 4b**) were all located in the middle of ICL3, indicating these conformational transitions may have potential roles in mediating the recruitment and interaction with kinases, and subsequent phosphorylation. The conformational heterogeneity is reduced but still exists when we further move the sensor toward intracellular end of TM5, as shown by the FRET distribution of 148/227, which exhibits a main high-FRET (∼0.95) population with a small tail of mid-FRET (∼0.65) population. Of interest, T227C was resolved in inactive and active structures, whereas structural comparison showed subtle conformational change in this region (**Extended data Fig. 5c**). It should be noted that the ICL3 was replaced with a BRIL protein to facilitate crystallization ^30^, thus, the conformation of TM5 observed in the inactive state maybe affected by protein engineering. In spite of the conformational heterogeneity, the high-FRET populations (ranging from 0.9 to 0.95) predominated the distribution for all sensors in ICL3 (**Fig. 4c, e and Extended data Fig. 5b**). This suggests that, in the apo state, the entire ICL3 prefers an overall compact conformation or a set of conformations with fast exchange rate (<100 ms) that are in close proximity to the intracellular surface of the TM domain (more specifically TM4).

### Negative allosteric regulation of G protein coupling by ICL3

The results presented above resemble our recent smFRET study on the disordered C-terminus (CT) of β2AR, where we observed a high-FRET distribution for a TM4-CT sensor in Apo state and showed that the β2AR-CT interacts with the intracellular surface of the receptor to directly inhibit G protein coupling ^31^. To investigate the role of ICL3 in α_2A_AR mediated G protein signaling, we made a α_2A_AR mutant in which the ICL3 was truncated (α_2A_AR△ICL3) (Fig. 4B). To directly compare the G protein coupling efficiency against the full-length α_2A_AR and α_2A_AR△ICL3, we utilized the TM4-TM6 sensor and titrated the effect of GoA by adding increasing amounts of GoA^GDP^ to immobilized α_2A_AR bound to Norepi (**Fig. 5a-b**). We observed a GoA^GDP^ concentration-dependent increase in the low-FRET active population, which plateaued at around 10 μM GoA^GDP^ (**Fig. 5c**). Interestingly, GoA^GDP^ exhibited different ability to shift the FRET population for different α_2A_AR constructs, as shown by the plotted concentration-response curve, the full-length α_2A_AR showed a ∼4 fold higher EC50 value than the α_2A_AR△ICL3, while maximum low-FRET occupancy was significantly lower than that of the α_2A_AR△ICL3 (**Fig. 5c**). In line with smFRET data, GTP-turnover assay also showed higher efficacy in Noerpi elicited GoA activation for α_2A_AR△ICL3, although we did not observe significant basal activity for either construct (**Fig. 5d**). These results suggest that the ICL3 of α_2A_AR functions as a negative allosteric regulator to attenuate agonist-stimulated GoA activation, possibly via two mechanisms: first, ICL3 may directly occupy the space for G protein binding like the β2AR-CT, as indicated by the smFRET data showing predominant high-FRET populations; second, as ICL3 is directly linked to TM6, the closed compact conformation of ICL3 may restrain the outward movement of TM6 necessary for activation, thereby stabilizing the receptor in a more inactive state. Interestingly, the negative allosteric regulation of ICL3 on receptor activity has also been reported in other GPCRs including β2AR, neuropeptide Y1, M3 muscarinic receptor and bitter taste receptors ^32–35^.

**Fig. 5.**
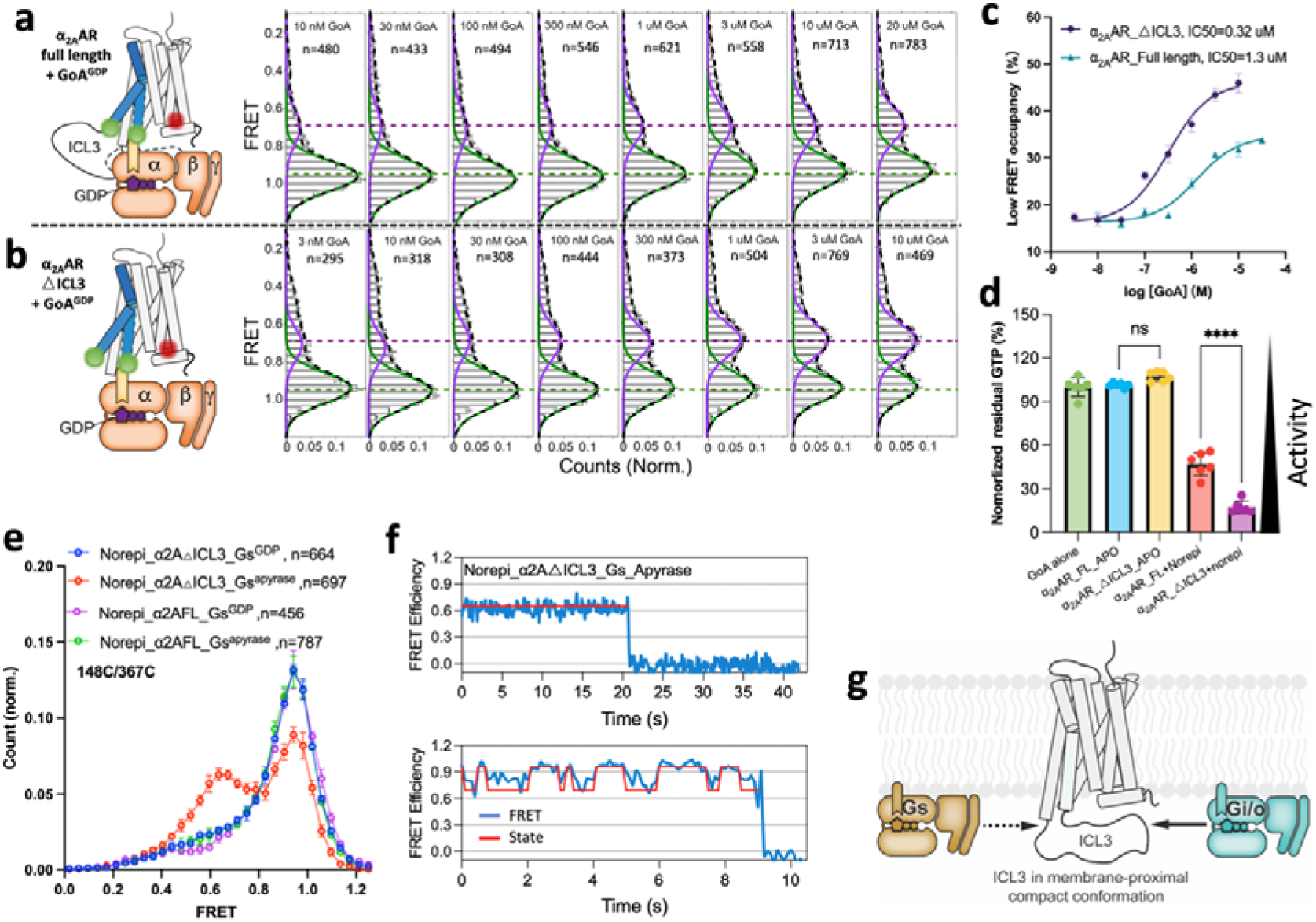
Negative allosteric regulation of G protein coupling by ICL3. **(a-b)** smFRET distributions of the TM4-TM6 sensor in the presence of Norepi and a range of GoA^GDP^ concentrations for the full length (**a**) and ILC3 truncated (**b**) α_2A_AR. Green and purple lines represent Gaussians fitted to the high-FRET inactive and low-FRET active states, respectively. Green and purple dash lines indicate the distinct mean FRET values. Black dashed line represents the cumulative fitted distributions. n represents the number of traces used to calculate the corresponding histograms. Data are mean ± s.e.m. from at least three groups of independent movies. Each group contains at least 3 movies. **(c)** GoA^GDP^ concentration-dependent curve of the TM4-TM6 sensor low-FRET state occupancy for the full length (cyan) and ILC3 truncated (purple) α_2A_AR. Data are mean ± s.e.m. **(d)** GTP turnover activity for the full-length and ILC3 truncated α_2A_AR. Data are mean ± s.e.m. of 6 replicates. **(e)** Overlaid raw smFRET histograms of the TM4-TM6 sensor in Norepi_Gs^GDP^ and Norepi_Gs^apyrase^ bound conditions for the full-length and ILC3 truncated α_2A_AR. **(f)** Representative single-molecule FRET time traces (blue) overlaid with predicted state sequence (red) showing Gs coupled low-FRET active state (upper) and transition between inactive and active states (bottom). **(g)** The membrane-proximal feature of ICL3 comtributes to the selective coupling of α_2A_AR with Gi/o over Gs.

In addition to Gi/o proteins, previous studies have shown that α_2A_AR can also weakly couple to Gs protein ^30,36^. Recent bioinformatics and functional studies showed that longer ICL3s (>45 amino acids) in GPCRs can regulate signaling specificity by inhibiting G protein subtypes that weakly couple to the receptor ^37,38^. To examine whether this also holds true for α_2A_AR, we investigated the effect of ICL3 on Gs coupling. We first imaged the TM4-TM6 sensor in the presence of Norepi and Gs^GDP^. However, we did not see an obvious increase of low-FRET population for both full-length α_2A_ARFL and α_2A_AR△ICL3 (**Fig. 5e**). We reasoned that the α_2A_AR-Gs complex might be very unstable in the presence of GDP, as also evidenced by the α_2A_AR-GoA^GDP^ complex, in which the low-FRET state occupancy was significantly lower than that of nucleotide-free complex (**Extended data Fig. 5d**). We then imaged the apyrase pretreated α_2A_AR-Gs complex. In this case, we observed a dramatic increase of the low-FRET active population for α_2A_AR△ICL3-Gs^apyrase^ (**Fig. 5e**). Inspection of individual trajectories also revealed similar stable low-FRET state or transitions between low and high-FRET states (**Fig. 5f**) as observed in the α_2A_AR-GoA^apyrase^ complex (**Fig. 1d**). In contrast, the full-length α_2A_AR-Gs^apyrase^ remains a dominant high-FRET peak (**Fig. 5e**), suggesting that the ICL3 indeed can inhibit Gs coupling. These data therefore reinforce the notion that the large ICL3 plays a role in fine-tuning the G protein coupling specificity (**Fig. 5g**).

### Modulation of ICL3 conformational dynamics by kinase phosphorylation

Recent biophysical studies on the C-terminal domains of several GPCRs (V2R, GHSR, and β2AR) have demonstrated that the structural dynamics of these disordered regions are strongly influenced by GRK phosphorylation or charge distribution^39^. Building on these findings, we investigated the impact of phosphorylation on the conformational dynamics of the ICL3 of α_2A_AR. To assess the effects of GRK-specific phosphorylation, we first dephosphorylated the purified receptor using lambda phosphatase (λpp) to remove basal phosphorylation present in the cellular environment. Consistent with prior biochemical studies in mammalian cells ^40^, we observed high basal phosphorylation of α_2A_AR in insect cells (**Extended data Fig. 6a**). Given that the ICL3 of α_2A_AR has been identified as an excellent substrate of GRK2 ^25^, we used GRK2 for in vitro phosphorylation assays. The dephosphorylated receptor can be efficiently phosphorylated by GRK2, reaching a plateau within approximately 2 hours (**Extended data Fig. 6b-c**).

We then imaged ICL3 FRET sensors under λpp-dephosphorylated and GRK2-phosphorylated conditions and observed pronounced effect of phosphorylation on the FRET distributions of these sensors (**Extended data Fig. 6d-g**). Upon dephosphorylation, the 148/322 (**Extended data Fig. 6e**) and 148/292 (**Extended data Fig. 6f**) pairs, located in the ICL3 middle region near the identified phosphorylation sites (**Fig. 4b**), exhibited the most dramatic shifts of the FRET population, from dominant high-FRET states to mid-FRET states with broader distributions. The FRET distributions of 148/352 (**Extended data Fig. 6d**) and 148/262 (**Extended data Fig. 6g**) sensors also become broader with increased mid to low FRET populations, indicating more heterogeneous and extended conformations. These findings suggest that phosphorylation plays a key role in stabilizing the high-FRET, compact conformations of ICL3. Supporting this notion, we observed an increase in the high-FRET populations for all sensors following phosphorylation by GRK2 (**Extended data Fig. 6d-g**). Notably, the FRET histograms for GRK2-phosphorylated samples differed from those of the basal-phosphorylated samples, likely reflecting distinct phosphorylation patterns.

In contrast to the FRET sensors in the ICL3 loop region, the phosphorylation state had minimal impact on the FRET distributions of TM4-TM5 (148/227) and TM4-TM6 (148/367) sensors (**Extended data Fig. 6h-i**), suggesting that the phosphorylation on ICL3 does not significantly affect TM5 and TM6 displacement. However, it remains possible that the conformational changes of ICL3 induced by phosphorylation could alter intra-molecular interactions in the transmembrane domain. Indeed, prior NMR studies on the M2R and β2AR have shown that GRK phosphorylation at intracellular loops can modulate the conformation of the transmembrane domain ^41,42^.

We next investigated whether phosphorylation-induced conformational changes in ICL3 contribute to the negative allosteric modulation of G protein coupling. To this end, we utilized the TM4-TM6 (148/367) sensor to image the dephosphorylated and GRK2 phosphorylated receptor in the presence of agonist and saturation concentration of GoA^GDP^ (20uM) to compare the efficacy of complex formation (**Extended data Fig. 6j**). Interestingly, we observed a slight, albeit not statistically significant, increase in active state occupancy for the dephosphorylated receptor compared to the basal- or GRK2-phosphorylated receptor (**Extended data Fig. 6k**). We then performed GTP-turnover assays and found that the dephosphorylated receptor showed significantly higher activity in activating GoA compared to phosphorylated receptor (**Extended data Fig. 6l**). Together, these results suggest that phosphorylation in ICL3 plays a critical role in stabilizing the compact conformation of ICL3 and contributes to the dampened G protein signaling. These findings provide a potential structural explanation for earlier functional and biochemical observations that the agonist-promoted desensitization of α_2A_AR was accompanied by phosphorylation in ICL3 ^12^.

### Activation dynamics of ICL3 modulated by ligand efficacy

The data present above suggest that full activation of α_2A_AR requires substantial conformational changes of ICL3 to create space for G protein coupling. To confirm this, we investigated the conformational dynamics of ICL3 of activated α_2A_AR. Native non-dephosphorylated receptor was used for this purpose to preserve the basal phosphorylation observed in mammalian cells ^40^. We first imaged the sensors in the presence of full agonist Norepi, but no obvious change in FRET distribution was observed, similar to that of TM6 (**Extended data Fig. 7**). Whereas incubation of the α_2A_AR with GoA in the absence of Norepi also showed negligible effect on the FRET distribution (**Fig. 6a, Extended data Fig. 7**), formation of the ternary complex with Norepi and GoA, upon treatment with apyrase, led to a dramatic shift of the FRET distribution for all sensors (**Fig. 6a**), indicating large conformational changes of ICL3 upon coupling to GoA. For the 148/352 and 148/322 sensors, we observed dramatic shifts to low-FRET states at ∼0.58 and ∼0.33, respectively, while the 148/292 and 148/262 sensors exhibited the emergence of a distinctive mid-FRET state (∼0.5) (**Fig. 6a**). The appeared new FRET states for these sensors indicate the stabilization of the G protein coupled active conformation of ICL3. Additionally, the TM4-TM5 (148C/227C) sensor showed increased occupancy of the mid-FRET state (∼0.65) (**Fig. 6a**), potentially representing a fully active conformation of TM5 not seen in previous structural studies.

**Fig. 6.**
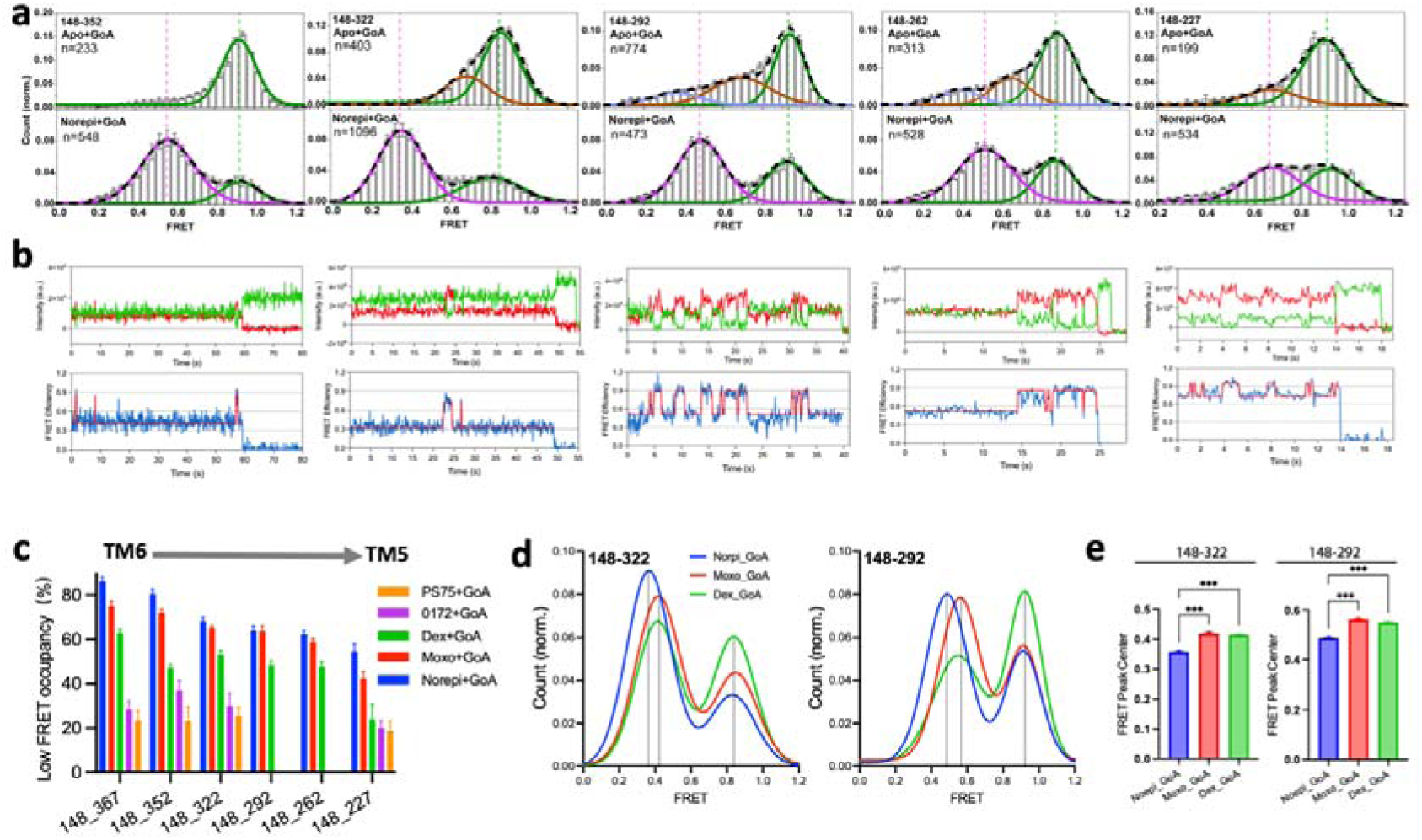
Conformational dynamics of ICL3 in active state. **(a)** smFRET distributions of TM4-ICL3 sensors in the Apo+GoA^apyrase^ (upper) and Norepi+GoA^apyrase^ (bottom) conditions with fitted Gaussians peaks. n represents the number of traces used to calculate the corresponding histograms. Data are mean ± s.e.m. from at least three groups of independent movies. Each group contains at least 3 movies. **(b)** Representative single-molecule fluorescence (donor, green; acceptor, red) and FRET time traces (blue) overlaid with predicted state sequence (red) showing two states transitioning for all TM4-ICL3 sensors in fully active α_2A_AR. **(c)** Comparison of low-FRET active state occupancy of different ligand conditions (incubated with GoA and apyrase) for TM4-TM6 sensor and TM4-ICL3 sensors. **(d)** Overlaid smFRET distributions (Gaussain Fitting) for 148/322 and 148/292 sensors in the presence of different agonists and GoA^apyrase^. **(e)** Comparison of low-FRET active state peak centers for 148_322 and 148_292 FRET sensors in different ligand conditions. Data are mean ± s.e.m. from at least three groups of independent movies.

As described above, the overall conformational landscape of ICL3 in the GoA-coupled activated α_2A_AR differed markedly from that of the apo state receptor (**Fig. 6a-b**). It’s possible that the ICL3 forms direct interactions with GoA and/or becomes partially ordered upon G protein coupling. Notably, a recent HDX-MS study on the M3 muscarinic receptor revealed interactions between the large ICL3 and Gq protein, which dramatically altered its dynamics ^35^. Indeed, in contrast to the apo state, the FRET distributions for all ICL3 sensors displayed a similar bimodal shape in the active state (**Fig. 6a**), exhibiting more concerted conformational dynamics across the entire loop. A two-state model was also revealed for all sensors from individual trajectories (**Fig. 6b**), similar to that of the TM4-TM6 sensor (**Fig. 1d**). These observations indicate that the conformational changes of ICL3 may be coupled with TM6 outward motions upon G protein coupling. Notably, the occupancy of the low-FRET active state appeared to decrease as the FRET sensor was moved from the C-terminal of ICL3 (TM6) to the N-terminal (TM5) (**Fig. 6c**), suggesting looser coupling between TM6 and the N-terminal part of ICL3.

To further investigate the potential role of ICL3 in ligand-specific signaling, we imaged each of the ICL3 FRET sensors in the presence of different ligands and GoA (**Extended data Fig. 8**). Similar to the TM4-TM6 sensor, ligands with different efficacies exhibited varying ability to stabilize the active conformation of ICL3, as evidenced by the distinct low-FRET occupancies (**Fig. 6c, Extended data Fig. 8**). For the 148/352 and 148/322 sensors, we observed almost the same order (Norepi > Moxo > Dex > 0172 and PS75) of low-FRET state occupancy as observed for the TM4-TM6 sensor for different ligands (**Fig. 6c, Extended data Fig. 8**), exhibiting positive correlation with ligand efficacies. Similar to Norepi, all other ligands seem to display a smaller structural change when moving the sensors toward the N-terminal region of ICL3 (**Fig. 6c, Extended data Fig. 8**). Of note, there was no obvious change in the FRET peak for the low efficacy G protein biased agonists 0172 and PS75 in the 148/292 and 148/262 sensors (**Extended data Fig. 8**). Also, there was no significant increase in the occupancy of the low-FRET active state in the 148/227 sensor for 0172 and PS75 (**Fig. 6c, Extended data Fig. 8**), suggesting that these weak biased agonists are less efficient in stabilizing conformational changes at the N-terminus of ICL3. Moreover, the full agonist Norepi stabilized active conformation in the middle region of ICL3 appeared to be distinctive from that stabilized by other ligands, as evidenced by the small but significant shift in the FRET peak center of the active state in 148/322 and 148/292 sensors (**Fig. 6d-e**). Notably, the observation that the full agonist Norepi stabilized the most pronounced conformational changes of ICL3 is in agreement with the orthosteric binding modes showing that Norepi is the only agonist forming hydrogen bond interactions with both TM5 and TM6 at the orthosteric pocket (**Extended data Fig. 9**). These findings align with previous live-cell ensemble FRET studies showing that the N-terminal and middle region of ICL3 respond less to weak partial agonists than to Norepi in terms of both amplitudes and kinetics ^43^. Taken together, these data reveal the allosteric coupling from the extracellular ligand-binding pocket to the distal ICL3, where different functional classes of ligands have distinct effects on the conformational states and dynamics of ICL3, in particular at the middle and N-terminal regions, therefore underscoring the significant role of ICL3 in determining ligand-specific functional outcomes.

## Discussion

An in-depth characterization of the conformational dynamics of GPCRs is important for understanding their functional diversity and complexity and may help in the rational design of highly specific drugs with minimal side effects. Previous structural and biophysical studies have been primarily restricted to the structured transmembrane domains ^44^, leaving a gap in our knowledge about the structure and dynamics of several intrinsically disordered and highly flexible regions. Single-molecule FRET imaging has been shown to provide valuable insights into the structure and dynamics of IDRs ^45,46^. Here, our smFRET studies offer a detailed characterization of real-time conformational changes in both the canonical TM6 and two structurally elusive loops, ECL2 and ICL3, during α_2A_AR activation. By integrating insights from both the structured and disordered regions of α_2A_AR, we present a holistic understanding of the molecular mechanism underlying GPCR signaling and regulation.

Decades of structural and biophysical studies of GPCRs have led to the widely accepted view that agonist binding can induce the outward displacement of TM6, creating an intracellular cavity for transducer binding. Rhodopsin is unique among GPCRs in that agonist alone stabilizes the fully outward conformation of TM6 ^47^. While for non-rhodopsin receptors, like β2AR, the rates or amplitudes of TM6 outward motions often correlate with ligand efficacy ^17^. Remarkably, our smFRET and DEER studies reveal that, for α_2A_AR, there is no obvious TM6 outward displacement upon agonist binding, whereas the active outward conformation is only captured in the presence of G protein (**Fig. 2**). Interestingly, a recent spectroscopic study of the μOR also revealed that TM6 displacement was not necessary for activation by most agonists other than two super agonists. Instead, agonist-specific receptor activation may rely on conformational changes in ICL2 rather than TM6 ^18^. Moreover, a recent simulation study of FFAR1 revealed a non-canonical activation mechanism driven by conformational changes of ICL2 rather than transmembrane helices ^48^. These findings reveal the complex conformational dynamics across the GPCR family, where different receptors have different activation mechanisms or energy landscapes in response to different ligands ^49,50^. Future studies might generate super agonists of α_2A_AR capable of overcoming the high energy barrier for TM6 outward movement in the absence of G protein.

Through smFRET, we are also able to delineate the conformational transitions of the partically disordered ECL2 of α_2A_AR between different conformational states in the absence of agonist and reveal that different functional classes of agonists can stabilize distinct conformational dynamics of ECL2. These findings highlight a common role of ECL2 in regulating the ligand efficacy and bias for GPCRs in spite of the sequence divergence ^27,51^. Cryo-EM structures have shown that current α_2A_AR agonists all bind to the small orthosteric pocket with similar poses (**Extended data Fig. 9**), and there is no reported allosteric ligand for α_2A_AR to date. Future drug development targeting the ECL2 of α_2A_AR may facilitate the discovery of small molecule drugs with better functional selectivity and therapeutic effects. Notably, there are already successful cases for the rational design of pharmacologically biased ligands by directly targeting ECL2 in other aminergic GPCRs ^52^.

α_2A_AR is one of the important GPCRs that has an exceptionally long ICL3 with unknown structural information ^6,12,37^. This study provides a comprehensive biophysical mapping of the conformational landscapes of this large ICL3 in different functional states at residue-level resolution. Our smFRET data, combined with functional analysis, reveal that ICL3 of α_2A_AR plays a pivotal role in fine-tuning signaling specificity through negative allosteric regulation of G protein coupling. In α_2A_AR, this effect arises primarily from the compact conformations of ICL3, which are situated close to the intracellular surface of TM domain, occupying the space required for G protein binding (**Fig. 4**–**5**). More importantly, we demonstrated how GRK2 phosphorylation modulates the conformational dynamics of ICL3, suggesting a mechanistic link between post-translational regulation and receptor function.

It is noteworthy that in the recent ensemble FRET study of β2AR ICL3, a low-FRET open state was observed in the Apo (inactive) state while agonist binding can stabilize the high-FRET closed state ^37^. These observations are in contrast to our smFRET data in α_2A_AR, indicating different ICL3 dynamics. Since ICL3 is directly linked to TM6, this may also reflect distinct TM6 dynamics between β2AR and α_2A_AR. Alternatively, these differences could arise from the varying labeling strategies employed in the studies.

Numerous biophysical studies have demonstrated ligand-specific conformations of GPCRs’ TM domain ^53–57^, but it remains unclear whether such differences extended to intracellular loops. Our smFRET analysis revealed distinct active conformations and dynamics in the middle region and N-terminus of ICL3 when bound to functionally different agonists in the presence of GoA. These findings suggest a critical role of ICL3 conformational dynamics in fine-tuning ligand efficacy, and that the signaling profile of an agonist cannot be solely determined by TM domain structural dynamics; significant contributions also come from the disordered regions. Further investigations are required to validate whether this mechanism is conserved across GPCRs with long ICL3 (i.e. muscarinic receptors and serotonine receptors).

In addition to the canonical signaling proteins (G proteins, GRKs and arrestins), the ICL3 of α_2A_AR has been reported to directly interact with several other cytosolic scaffolding proteins such as 14-3-3ζ and spinophilin to regulate arrestin binding ^58^. The observation that the middle region of ICL3 is capable of transitioning to a distal low-FRET state (**Fig. 4**) might be important for tethering these cytosolic proteins. This supports the notion that the loop acts as a “fishing line” ^6^, expanding the capture radius for interactions with cytosolic partners. Moreover, the ICL3 conformation appears to be of high plasticity, as evidenced by the re-shaped FRET distributions by phosphorylation or G protein coupling. Such conformational plasticity may allow ICL3 to interact with diverse cytosolic proteins. Moreover, our smFRET study on ICL3 offer clues to a mechanistic understanding of other unresolved functional properties of α_2A_AR. For example, previous studies revealed that membrane potential can modulate α_2A_AR signaling ^59^. A change of membrane potential may affect the charge distribution of ICL3 therefore stabilizing the loop into different conformational states and consequently resulting in distinct signaling behavior.

Taken together, this integrative study not only uncovers the complex signaling mechanisms encoded in both the structured TMD and IDRs of α_2A_AR, but also highlights that the dynamic plasticity of IDRs serves as a universal regulatory hub for deciphering receptor functional diversity. Along with the recent advance in de nove design of IDR binders ^60^, these insights would pave the way for developing next-generation precision therapeutics that target these dynamic IDRs, with potential implications extending to other GPCRs or membrane proteins featuring similar disordered regions.

## Acknowledgments

We thank Prof. Brian K. Schoichet (UCSF) for providing the α_2A_AR small molecules (PS75 and 0172), and Prof. Richard Lewis (School of Medicine, Stanford) for the use of their TIRF microscope. We thank Dr. John Janetzko for the help with smFRET data collection of AF555/Cy7 labeled samples and Dr. Betsy White for help with G protein preparations. This work was supported by the National Institutes of Health under award number R01GM135581 (M.T.L.), S10 OD025260 (M.T.L.) and R35NS137408 (B.K.K.). Jun Xu was supported by a Dean’s fellowship of Stanford Medicine School. P.G. is supported by the Deutsche Forschungsgemeinschaft (DFG, German Research Foundation) grant GRK GM 13/15.

## Author contributions

J.X. designed and validated constructs of the α_2A_AR, purified all the proteins, performed de-phosphorylation and phosphorylation of the α_2A_AR, performed GTP turnover assays, labelled the α_2A_AR for smFRET and DEER studies, collected and analyzed smFRET data. R.Q collected, procesed and analyzed the smFRET data. A.G. performed DEER experiments and data analysis. H.H. and D.W. performed cell signalling assays and radioligand binding assays under the supervision of P.G.. X.Y. performed the molecular docking studies. M.L. supervised DEER data collection and analysis. B.K.K. supervised the overall project. J.X. and B.K.K wrote the manuscript with contributions from all authors.

## Data and materials availability

All data and materials used in the analysis are available upon request.

## Supplementary Materials

Materials and Methods

References

Extended data Fig. 1 to 9

