## Supplementary figure for "The role of intrinsically disordered domains in regulating G protein coupled receptor signaling"

**The PDF file includes:**

Materials and Methods

References

Extended data Fig. 1 to 9

**Materials and Methods**

**α_2A_AR expression and purification**

The wild-type human α_2A_AR with an N-terminal HA signal sequence followed by a Flag tag and a C-terminal 8×His tag was cloned in the pFastBac1vector. The minimal-cysteine construct (α_2A_AR∆3) was created by introducing the mutations C38A, C401S, C442A into the wild-type α_2A_AR. Double-cysteine mutation constructs (TM4/TM6, TM4/ECL2 and TM4/ICL3) for smFRET and DEER experiments were generated based on the α_2A_AR∆3 construct.

Recombinant baculovirus for insect cell expression was made using the Bac-to-Bac system. Sf9 cells were grown in ESF921 insect cell Medium and were infected with recombinant baculovirus containing the α_2A_AR gene at a density of 4 × 10^6^ cells ml^−1^ in the presence of 5 μM rauwolscine. After 48 hours of infection at 27°C, the cells were spun down and cell pellets were stored at −80°C until use.

Thawed cell pellets were resuspended in hypotonic lysis buffer [10 mM tris, 1 mM EDTA, 5 μM rauwolscine, leupeptin (2.5 μg ml^−1^), and benzamidine (160 μg ml^−1^)]. Cell membranes were then spun down and solubilized with a buffer consisting of 20 mM Hepes (pH 7.5), 500 mM NaCl, 1% n-Dodecyl-b-D-Maltopyranoside (DDM), 0.2% sodium cholate, 0.03% Cholesteryl Hemisuccinate (CHS), 5 μM rauwolscine, leupeptin (2.5 μg ml^−1^), and benzamidine (160 μg ml^−1^). Nickel-NTA sepharose was added to the solubilized receptor and rotated for 2 hours at 4°C. The resin was spun down and washed in batch for three times with a buffer composed of 20 mM Hepes (pH 7.5), 500 mM NaCl, 0.1% DDM, 0.02% sodium cholate, 0.03% CHS, 5 μM rauwolscine, leupeptin (2.5 μg ml−1), and benzamidine (160 μg ml^−1^). The washed resin was poured into a glass column, and the receptor was eluted in the wash buffer supplemented with 250 mM imidazole.

The Ni-NTA chromatography–purified receptor was immobilized using anti-flag M1 affinity resin and was extensively washed with a buffer containing 20 mM Hepes (pH 7.5), 500 mM NaCl, 0.1% DDM, 0.02% sodium cholate, 0.003% CHS, 2 mM CaCl_2_. The receptor was then gradually exchanged into a buffer containing 20 mM HEPES pH 7.5, 100 mM NaCl, 0.01% lauryl maltose neopentyl glycol (MNG), 0.003% CHS supplemented with 2 mM CaCl_2_, and then eluted with same buffer containing 0.2 mg mL^−1^ flag peptide and 5 mM EDTA. The eluted receptor (2 to 3 ml) was concentrated to 500 μl using a 50-kDa molecular weight cutoff Millipore concentrator. The flag affinity chromatography purified receptor was then subjected to different biophysical and bichemical experiments.

**GoA heterotrimer expression and purification**

The GoA protein, which represents the heterotrimeric complex of the guanine nucleotide–binding protein Go subunit α (Gαo), Gβ, and Gγ subunits, was expressed and purified as previously described (*1*). In brief, human Gαo was cloned into pFastBac vector, Gβ1 with 3C protease-cleavable 6xHis-tag, and Gγ2 was cloned into pFastBac_Dual vector. The GoA was expressed in HighFive insect cells grown in ESF921 Medium. Cells were grown to a density of 3 million per milliliter and infected with Gαo and Gβ1γ2 baculovirus at a ratio of 10 to 20 ml liter^−1^ and 1 to 2 ml liter^−1^, respectively. After 48 hours of incubation, the infected cells were harvested by centrifugation and stored at −80°C until use.

Thawed cell pellets were resuspended in lysis buffer [10 mM tris (pH 7.5), 0.1 mM MgCl_2_, 5 mM β-mercaptoethanol (β-ME), 10 μM guanosine diphosphate (GDP), leupeptin (2.5 mg ml^−1^), and benzamidine (160 mg ml^−1^)] and stirred at room temperature for 15 min. Cell membranes were spun down and resuspended with solubilization buffer [20 mM Hepes (pH 7.5), 100 mM NaCl, 1% sodium cholate, 0.05% MNG, 5 mM MgCl2, 2 ml CIP, 5 mM β-ME, 15 mM imidazole, 10 μM GDP, leupeptin (2.5 mg ml^−1^), and benzamidine (160 mg ml^−1^)] using a Dounce homogenizer. The samples were stirred at 4°C for 40 min and then centrifuged for 30 min to remove insoluble debris.

Ni-NTA resin preequilibrated in solubilization buffer was added to the supernatant and shaken for 2 hours at 4°C. After incubation, the Ni-NTA resin was spun down and poured into a glass column and washed with 50 ml of solubilization buffer. The heterotrimeric GoA was gradually exchanged into E2 buffer [20 mM Hepes (pH 7.5), 50 mM NaCl, 0.2% MNG, 1 mM MgCl2, 5 mM β-ME, 10 μM GDP, leupeptin (2.5 mg ml^−1^), and benzamidine (160 mg ml^−1^)]. The protein was then eluted with E2 buffer supplemented with 250 mM imidazole.

The sample was then dephosphorylated by treating with 5 μl lambda phosphatase (supplemented with 1 mM MnCl_2_ for activity; New England Biolabs) and incubated at 4°C for 30 min. The Ni-NTA chromatography–purified GoA was further purified with a MonoQ column (GE Healthcare). The peak fractions of the MonoQ chromatography were collected and exchanged to E3 buffer [20 mM Hepes (pH 7.5), 100 mM NaCl, 0.05% MNG, 1 mM MgCl2, 10 μM GDP, and 50 μM Tirs (2-carboxyethy) phosphine (TCEP)] by repeated concentration and dilution using a 50-kDa molecular weight cutoff Millipore concentrator. The concentrated heterotrimeric GoA was aliquoted, flash frozen in liquid nitrogen, and stored at −80°C before use.

**Purification of GRK2**

GRK2 was expressed and purified as previously described (*2*). In brief, bovine GRK2 with 6xHis-tag was cloned into pFastBac and expressed in SF9 insect cells. SF9 cells were grown to a density of 4 × 10^6^ cells mL^−1^ and then infected with GRK2 baculovirus at a ratio of 10–20 ml L^−1^. After 48 h incubation at 27 °C, the infected cells were harvested by centrifugation and stored at −80 °C until use. Cell pellets were resuspended lysis buffer composed of 20 mM HEPES, 250 mM NaCl, 1 mM β-ME, 0.02% Triton-X100, 2.5 μg mL^−1^ leupeptin and 160 μg mL^−1^ benzamidine and were stirred at RT for 15 min. Cells were then broken by sonication for 5 min on ice-water bath. Cell debris was removed by centrifugation at 18,000×rpm and the supernatant was collected. Ni-NTA resin pre-equilibrated in lysis buffer were added to the supernatant and shake for 1.5 h at 4 °C. After incubation, the Ni-NTA resin was spun down and poured into a glass column, and then washed with 50 mL lysis buffer and then eluted with lysis buffer containing 250 mM imidazole. The Ni-NTA purified GRK2 was finally purified by SEC against 20 mM HEPES pH 7.5, 100 mM NaCl. The monodisperse peak fractions was collected and then concentrated using a 50 kDa molecular weight cutoff Millipore concentrator. The concentrated GRK2 was aliquoted, flash frozen in liquid nitrogen and frozen at −80 °C until use.

**Dephosphorylation of α_2A_AR**

To prepare dephosphorylated α_2A_AR, the flag purified receptor was diluted to 5 µM in 200 µl of dephosphorylation buffer (20 mM HEPES pH 7.5, 100 mM NaCl, 0.01% LMNG, 0.001% CHS, 2 mM MnCl_2_). Lambda phosphatase was added at 1:1000 (v/v, NEB) to initiate the reaction. After incubation at 22 °C for 30 min, the dephosphorylated receptor was purified by SEC against 20 mM HEPES pH 7.5, 100 mM NaCl, 0.01% LMNG, 0.001% CHS to remove excess lambda phosphatase. Monodisperse peak fractions containing α_2A_AR were pooled and subjected to fluorophore labeling or GRK2 phosphorylation. The extent of dephosphorylation was evaluated by Pro-Q (sigma) and Coomassie blue staining.

**Phosphorylation of α_2A_AR**

Excess purified GRK2 was mixed with the dephosphorylated α_2A_AR in a buffer composed of 20 mM HEPES pH 7.5, 100 mM NaCl, 0.01% MNG, 0.003% CHS, 2 mM MgCl2, 500 μM ATP and 500 μM NorEpi to initiate the phosphorylation at room temperature. The extent of phosphorylation at various time points (from 0 to 240 min) was evaluated by Pro-Q and Coomassie blue staining. The phosphorylated receptor was further purified with anti-flag chromatography and SEC to remove excess GRK2 and agonist NorEpi. Monodisperse peak fractions containing α_2A_AR were pooled and subjected to fluorophore labeling.

**Preparation of α_2A_AR labeled with fluorophores**

α_2A_AR∆3 with introduced cysteine mutations was labelled by commercial maleimide-conjugated Alexa Fluor 555 (AF555) and Alexa Fluor 647(AF647) or by maleimide-conjugated AF555 and sulfo-Cy7, respectively. Flag purified α_2A_AR with basal-phosphorylation, or dephosphorylated by lambda phosphatase, or phosphorylated with GRK2, was diluted to 3 µM in 200 µl of labelling buffer (20 mM HEPES pH 7.5, 100 mM NaCl, 0.01% LMNG, 0.001% CHS). 5 µM of donor fluorophore and 10 µM of acceptor fluorophore were added into the reaction. After incubation at 22 °C for 20 min, free dyes were quenched with 5 mM L-cysteine. The labeled receptor was then purified by SEC against 20 mM HEPES pH 7.5, 100 mM NaCl, 0.01% LMNG, 0.001% CHS. Monodisperse peak fractions containing α_2A_AR were pooled, aliquoted, flash frozen with 15% glycerol and stored at −80°C before use.

**Preparation of α_2A_AR labeling with nitroxide spin label**

To prepare samples of the α_2A_AR for DEER studies, flag purified α_2A_AR∆3 with TM4-TM6 cysteine mutations (148C/367C) was diluted to 10 µM in labelling buffer (20 mM HEPES pH 7.5, 100 mM NaCl, 0.01% LMNG, 0.001% CHS). Nitroxide spin label reagent 3-(2-iodoacetamide)-proxyl (IAP) was added to a final concentration of 100 µM. After incubation at room temperature for 45min, another 100 µM of IAP was added to the reaction to make a final concentration of 200 µM. After incubation at room temperature for another 45min, the reaction was quenched with 5 mM L-cysteine and was injected into an SD200 increase 10/300 column equilibrated with SEC buffer (20 mM HEPES pH 7.5, 100 mM NaCl, 0.01% LMNG, 0.001% CHS in D2O). Fractions of the monodisperse peak were pooled and equally divided. Ligands were added to each tube at a final concentration of 1 mM. One tube of protein was kept without ligand. The α_2A_AR and ligand were incubated at room temperature for 2 h. Protein in each individual tube was mixed with 15% (v/v) D8-glycerol and flash frozen. NorEpi-bound receptor was prepared in two copies, one of them was mixed with a twofold molar excess of GoA and incubated for 60 min at room temperature. 1:100 apyrase (v/v, NEB) was added to the G-protein samples to remove free GDP and incubated for 1 h at room temperature.

**GTP turnover assay**

The α_2A_AR, including full length receptor with basal-phosphorylation, dephosphorylation or GRK2 phosphorylation, as well as ICL3 truncated receptor, for GTPase-GloTM assay were expressed and purified as described above and frozen at −80°C before use. The GTPase reaction was initiated by mixing GoA and α_2A_AR in 5 μL reaction buffer (20 mM HEPEs, 100 mM NaCl, 0.02% MNG, 1 mM MgCl_2_, 10 μM GTP, 10 μM GDP, with or without 500 μM NorEpi) in a 384-well plate. GoA and α_2A_AR were fixed at a final concentration of 0.5 μM and 1 μM, respectively, in the reaction system. For every independent experiment, GoA alone was set as a reference. The GTPase reaction was incubated at room temperature (22-25°C) for 1 h. After incubation, 5 μL reconstituted 1xGTPase-GloTM Reagent (Promega) was added to the completed GTPase reaction, mixed briefly and incubated with shaking for 30 minutes at room temperature (22-25°C) to convert the remaining GTP into ATP. Then 10 μL Detection Reagent (Promega) was added to the system and incubated in the 384-well plate for 5-10 minutes at room temperature (22-25°C) to convert the ATP into luminescent signals. Luminescence intensity was quantified using a Multimode Plate Reader (PerkinElmer) luminescence counter. Data were normalized to GoA alone (Emax %) and analyzed using GraphPad Prism 10.

**Single-molecule FRET experiments and analysis**

All smFRET experiments were performed at room temperature (22 °C) following previous protocol with some modifications (*3*). For all Alexa Fluor 555 and Alexa Fluor 647 labeled samples, single-molecule FRET recordings were performed using a home-built through-the-objective total internal reflection fluorescence (TIRF) imaging system based on a Zeiss Axiovert S100 TV microscope equipped with a Fluar 100 × 1.45 NA oil-immersion objective (Zeiss). Alexa Fluor 555 (donor) and Alexa Fluor 647 (acceptor) fluorophores were excited by 532 and 637 nm wavelength lasers (OBIS 532 nm LS 150 mW, Coherent and OBIS 637 nm LX 140 mW, Coherent). Donor and acceptor emission were separated by a 652 nm dichroic beamsplitter (Semrock) and passed through 580/60 nm and 731/137 nm bandpass filters (Semrock) mounted in an OptoSplit-II beamsplitter (Cairn Research), projecting the images side-by-side onto an EM-CCD camera (iXon DU897E, Andor). Hardware and image acquisition were controlled by BeanShell scripts in μManager, and image sequences were stored as stacked TIFF files.

The α_2A_AR was immobilized on the cover slip via biotinylated M1 Fab and neutravidin. In brief, the assembled glass chamber, which had been cleaned and passivated with biotin-polyethylene glycol (PEG), was incubated with 0.2 mg ml^−1^ neutravidin in wash buffer 20 mM HEPES 7.5, 100 mM NaCl. One minute later, the unbound neutravidin was flushed out with the wash buffer. Then, 25 nM biotinylated M1 Fab was incubated in the channel for one minute and the unbound M1 Fab was flushed out by wash buffer. The N-terminal Flag-tagged, fluorophore-labelled α_2A_AR was diluted to around 20 nM in incubation buffer (20 mM HEPES pH 7.5, 100 mM NaCl, 0.01% LMNG, 0.001% CHS, 2 mM CaCl_2_, 5 mM MgCl_2_ and 100 µM ligand) and incubated on ice for 1 h before measurement. The α_2A_AR was diluted to about 300-500 pM and injected into the chamber. The unbound α_2A_AR was removed by imaging buffer (incubation buffer + 100 µM cyclooctatetraene (COT), + 1% D-glucose, 1 mg ml^-1^ glucose oxidase and 0.04 mg ml^-1^ catalase). Movies were taken at a frame rate of 10 s^−1^ using the BeanShell scripts in μManager. For measurement in complex with GDP-free GoA, 20 nM α_2A_AR in the presence of 100 µM ligand was incubated with 15 µM GoA for 60 min followed by addition of 1:100 (v/v, NEB) apyrase. After incubation on ice for overnight at 4°C, the complex was diluted and injected into the chamber and measured following the same protocol above.

The movies were processed and analyzed using previously released custom Python scripts (*3*). In brief, traces were extracted and selected for final analysis if all of the following criteria were met: (1) the signal-to-noise ratio was ≥5; (2) the acceptor bleached in a single step before the donor bleached; (3) the γ factor was between 0.5 and 2.5; (4) spontaneous fluctuations of donor and acceptor fluorescence were negatively correlated; and (5) if a donor bleach event was recorded, it occurred in a single step. The FRET ratio E was calculated at each time point prior to bleaching as E = IA/(IA + γID), where IA and ID are the respective acceptor and donor fluorescence values. γ correction was used to correct for differences in photon emission and detection probabilities of donor and acceptor fluorophores. The value of γ was empirically determined for each individual molecule, and the correction was applied by multiplying the donor fluorescence signal by γ before calculating the smFRET ratio. A FRET histogram was constructed for each trace by distributing FRET amplitudes into 30 bins in a [–0.25, 1.25] range. Histograms were normalized by dividing each bin by the total number of FRET points in the trace. An ensemble histogram for multiple molecules was constructed by summing the normalized histograms of the individual traces and dividing each bin by the total number of molecules. Occupancies of high-FRET and low-FRET states observed in the histograms were calculated using Gaussian Fitting in Origin software. The transition states of fluorescence traces were parameterized by fitting a Hidden Markov model (HMM). The cumulative frequency count of low-FRET dwell times for each condition was fitted in Origin software to single exponential decay curves.

For samples labeled with Cy3 and Cy7, data were acquired using a lab-built prism-based TIRF microscope. Photons emitted from Cy3 and Cy7 were collected through a 1.27 NA 60x water-immersion objective (Nikon) and split onto 596 two cooled EMCCD cameras (Andor iXon Ultra 888) connected by a dichroic mirror for Cy3/Cy7 pair housed in a TwinCamTM (Cairn). Movies were recorded with Andor Solis (i) software (Andor) at a 100 ms time resolution. The movies were processed and analyzed using SPARTAN3.7 with similar data criteria as Alexa Fluor 555 and Alexa Fluor 647 labeled samples.

**Molecular Docking**

Molecular docking was performed using Glide (Schrödinger, LLC, New York, NY, 2024) with the 7EJ8 structure of the α_2A_AR. The protein structure was prepared using the Protein Preparation Wizard in Maestro with default settings, including hydrogen addition, bond order assignment, protonation at pH 7.0 ± 2.0 using Epik, and restrained minimization. Ligands were prepared from SMILES format using LigPrep with the OPLS4 force field, generating possible ionization states at pH 7.0 ± 2.0 using Epik Classic. Desalting and tautomer generation were enabled. Stereoisomer generation was set to retain specified chiralities while varying other chiral centers, with a maximum of 32 stereoisomers per ligand. Receptor grid generation was centered on the ligand binding site using default parameters. Docking was performed using the Standard Precision (SP) mode of Glide, generating 5000 poses per ligand, with 400 best poses retained for energy minimization. The docking protocol included flexible ligand sampling with nitrogen inversion and ring conformation sampling. Epik state penalties were added to the docking score, and bias sampling was applied for predefined functional groups, penalizing nonplanar amide conformations. The top 10 docked poses per ligand were saved, and GlideScore was used to evaluate binding affinity, with more negative scores indicating stronger binding. The docked poses were analyzed for key binding interactions.

**DEER experiments and analysis**

α_2A_AR samples were prepared at a final concentration of ~50 μM and volume of 13 μL, loaded into borosilicate capillaries (1.4 mm ID, 1.7 mm OD; VitroCom), and flash-frozen in liquid nitrogen. DEER experiments were conducted as previously described (*4*) at Q-band (~33.68 GHz) using a Bruker Elexsys 580 spectrometer equipped with a SpinJet AWG, EN5107D2 resonator, variable-temperature cryogen-free cooling system (ColdEdge Technologies Inc.), and a 150 W TWT amplifier (Applied Systems Engineering Inc.). All measurements were performed at 50 K. Dipolar evolution data were acquired using a dead-time-free 4-pulse DEER sequence with gaussian pulses (*5*) and with 16-step phase cycling.

The experimental parameters used for DEER data collection were: π/2, π_obs_, and π_pump_ pulse lengths of 40 ns; a frequency offset (Δv) of 90 MHz; d1 = 250 ns; d2 = 5150 ns; shot repetition time = 2000 µs; shots per point = 4; and integration window = 40 ns. The optimal microwave power (i.e., pulse amplitude) for the π/2, π_obs_, and π_pump_ pulses were determined using transient nutation experiments, where pulse amplitudes were adjusted to maximize the inversion of the Hahn echo (*6*). Pump pulses were applied to the maximum intensity of the field swept echo detected absorption spectrum. Observe pulses were applied at a frequency 90 MHz lower than the pump pulses.

DEER data were processed using DeerAnalysis 2021 (*7*), which employs two fitting routines: neural network analysis (DEERNet (*8*), Spinach revision 5662) and Tikhonov regularization (DeerLab 0.9.1) (*9*). The consensus fit represents the mean of both methods, with reported 95% confidence intervals also incorporating errors from both methods. Time traces were normalized to signal intensity at t = 0, and distance distributions were area normalized. Custom Python scripts were used for plotting the dipolar evolution time traces and the distance distributions.

**Cell-based BRET assays**

The human wild type α_2A_AR and the respective mutants all carrying an N-terminal HA-signal sequence and a FLAG-tag were cloned to pcDNA3.1 for G protein activation and for β-arrestin-2 recruitment assays, respectively, using polymerase chain reaction and Gibson Assembly. These constructs were used for functional experiments applying BRET biosensor based assays. G protein activation was monitored with Gα_i1_-RLucII together with Gβ_1_ and Gγ_2_-GFP_10_ (*10, 11*). Assessment of arrestin recruitment was performed by enhanced bystander BRET using CAAX-rGFP and β-arrestin-2-RLucII as biosensors in the presence of GRK2(*12*). In brief, HEK293T cells (gift from the Chair of Physiology, FAU Erlangen-Nürnberg) were transfected with 200 ng receptor plasmid for G protein activation (receptor:Gα:Gβ:Gγ ratio 4:1:2:8) or 100 ng receptor plasmid for β-arrestin recruitment (receptor: β-arrestin:CAAX:GRK2 ratio 5:1:15:5) using linear polyethyleneimine (PEI, Polysciences, 3:1 PEI:DNA ratio). The DNA was complemented to a total amount of 1 µg DNA per 3x10^5^ cells with ssDNA (Sigma Aldrich) and 10,000 cells per well were transferred into 96-well half-area plates (Greiner, Frickenhausen, Germany). After 48 h the cell medium was exchanged with PBS (phosphate buffered saline) and cells were stimulated with ligands at 37°C for 10 min. Coelenterazine 400a (abcr GmbH, Karlsruhe, Germany) at a final concentration of 2.5 μM was added 5 min before measurement. BRET was monitored on a Clariostar plate reader (BMG, Ortenberg, Germany) with the appropriate filter sets (donor 410/80 nm, acceptor 515/30 nm) and was calculated as the ratio of acceptor emission to donor emission. BRET ratio was normalized to the effect of buffer (0%) and the maximum effect of norepinephrine (100%). For each compound 3 to 8 individual experiments were performed each done in duplicates.

**Radioligand binding assays**

Receptor binding affinities for the α_2A_AR receptor wild-type and mutant were determined as described previously (*10, 13*). In brief, membranes were prepared from HEK293T cells transiently transfected with the cDNA for human α_2A_AR wild-type or the mutant α_2A_AR∆3. Receptor densities (B_max_ value) and specific binding affinities (K_D_ value) for the radioligand [³H]RX82,1002 (specific activity 52 Ci/mmol, Novandi, Södertälje, Sweden) were determined as 1,000 fmol/mg protein and 0.50 nM for the wild-type, and 2,000 fmol/mg protein and 0.50 nM for the α_2A_AR∆3 mutant, respectively. Competition binding was performed by incubating membranes in binding buffer (50 mM TRIS at pH 7.4) at final protein concentrations of 6 μg/well with the radioligand (final concentration 0.5 nM) and varying concentrations of norepinephrine for 60 minutes at 37 °C. Non-specific binding was determined in the presence of unlabeled RX82,1002 at 10 μM. Cell-based determination of binding affinity was performed with HEK293T cells transiently transfected with α_2A_AR or α_2A_AR∆3. Two days after transfection, cells were harvested and diluted in binding buffer to a final number of cells of 2,000 cells/well and used for binding experiments as described above. The resulting competition curves were analyzed by nonlinear regression using the algorithms implemented in Prism 10.4.2 (GraphPad Software, San Diego, CA) to provide IC_50_ values, which were subsequently transformed into a K_i_ values applying the equation of Cheng and Prusoff (*14*). For binding experiments with homogenates mean K_i_ values (± SD) for norepinephrine were determined with 830 ± 230 nM for α_2A_AR and 1000 ± 420 nM for α_2A_AR∆3 derived from 5 single experiments each performed in triplicates while for whole cell binding K_i_ values were measured as 3600 ± 1100 nM for α_2A_AR (n=2) and 4900 ± 1300 nM for α_2A_AR∆3 (n=4).

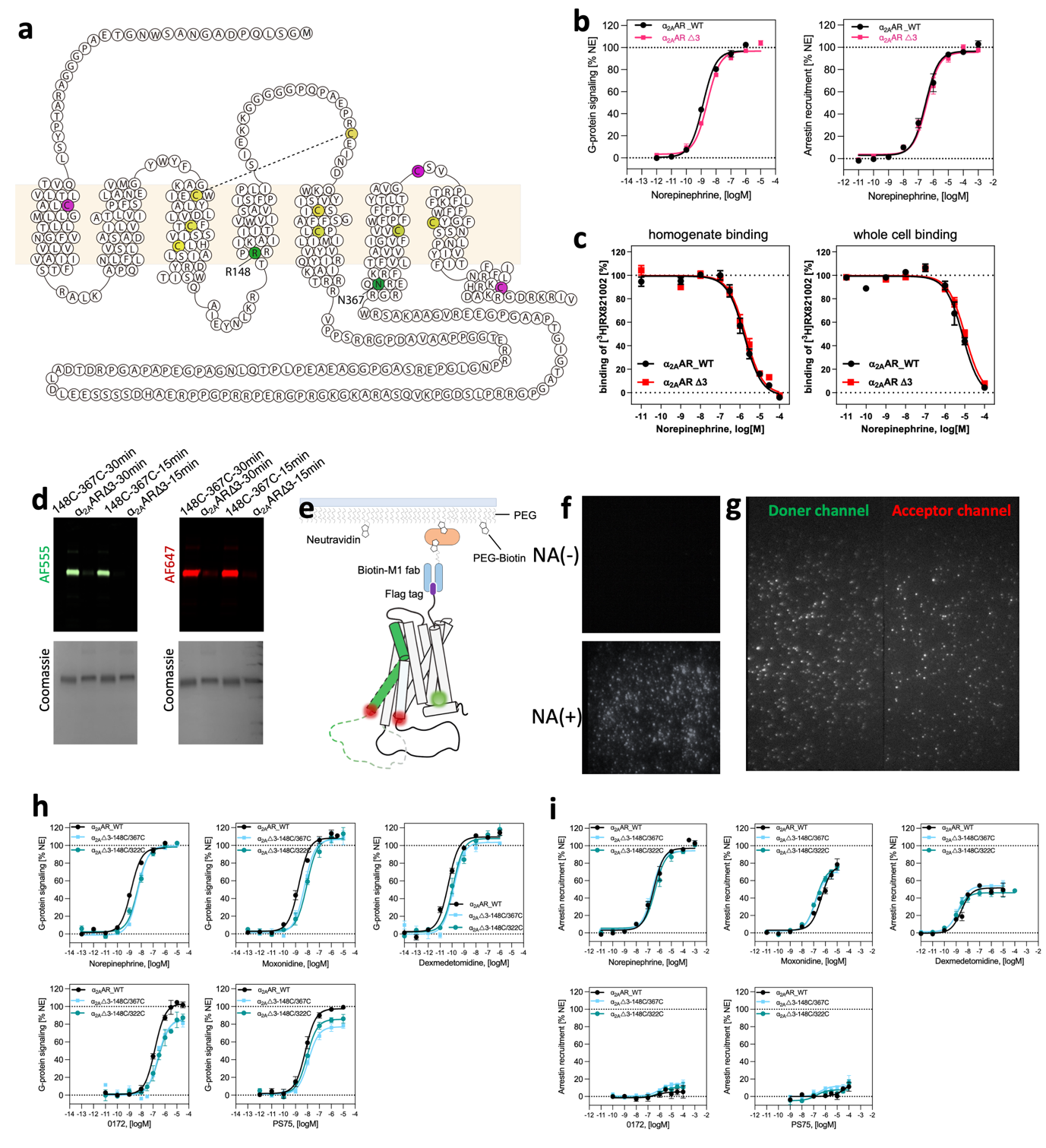

**Extended data Fig.1 Validation of smFRET construct.** (**a**) Design of the minimal cysteine construct of α_2A_AR. Mutated cysteine residues were shown in purple. Labeling sites at TM4 and TM6 were shown in green. Retained cysteine residues were shown in yellow. (**b-c**) Functional validation of α_2A_AR∆3 using cell-based functional assays (**b**) and radioligand binding assays (**c**). (**d**) Purified α_2A_AR∆3 background and with cysteines introduced (148C/ 367C) were analyzed with SDS-PAGE and imaged to detect AF555 fluorescence, AF647 fluorescence, and Coomassie stain. (**e**) Schematic of smFRET experiments for TM4-TM6 sensor (148/367) and receptor immobilization. (**f**) Specificity of neutrAvidin-mediated receptor immobilization. Frame capture from immobilization movies showing labelled α_2A_AR -148C/367C on neutrAvidin-free (−NA) or neutrAvidin-coated (+NA) surfaces. (**g**) Frame capture from representative movies showing doner and acceptor channel. (**h-i**) Functional validation of TM4-TM6 and TM4-ICL3 sensors for five selected agonists in G protein (**h**) and arrestin (**i**) pathways.

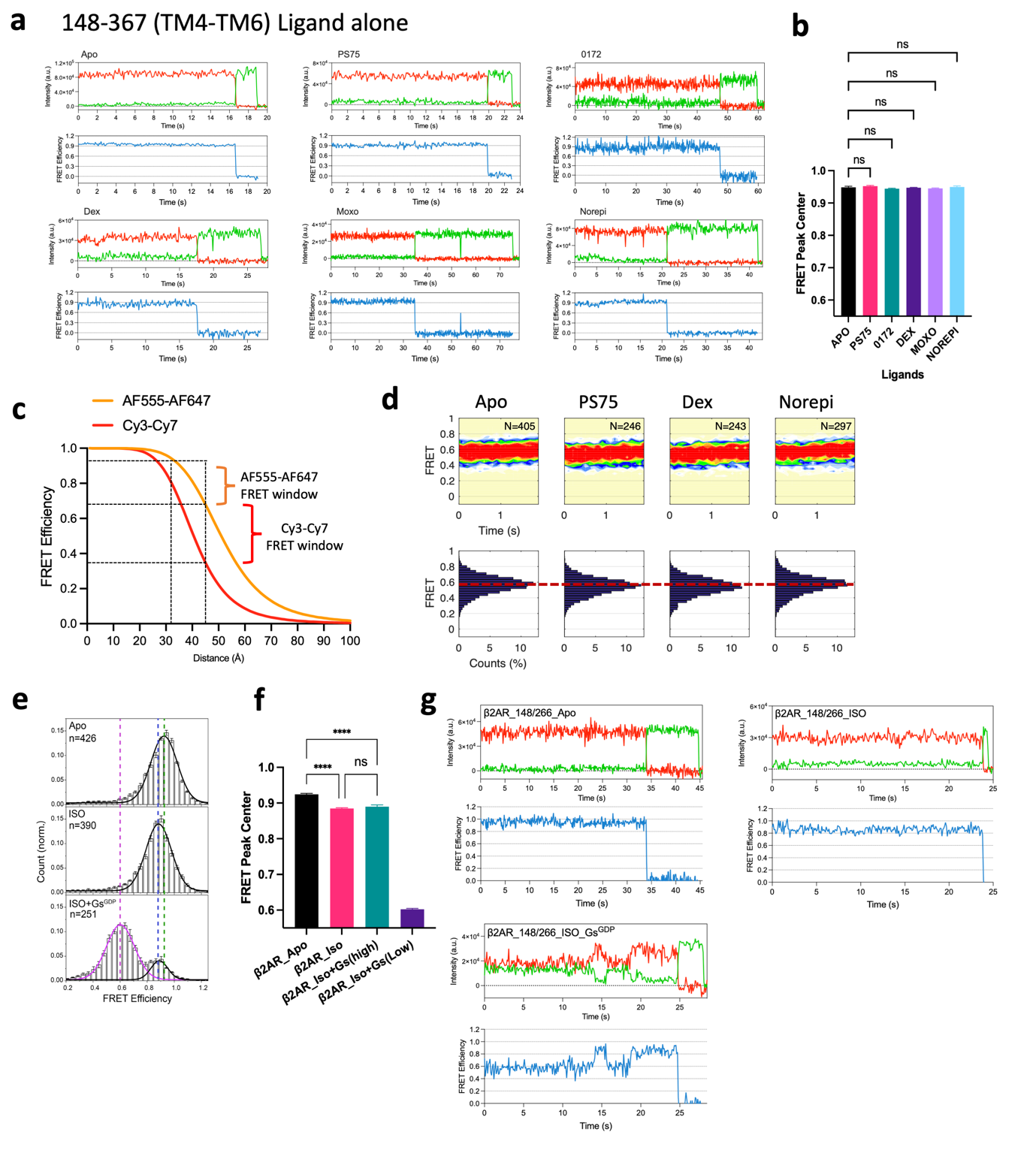
**Extended data Fig.2 TM6 conformational dynamics of α_2A_AR and β2AR.** (**a**) Example fluorescence (green for AF555; red for AF647) and FRET (blue) time traces for α_2A_AR -148C/367C imaged in the absence (Apo) and presence of saturating ligands. (**b**) Comparison of FRET peak centers for α_2A_AR -148C/367C in the absence (Apo) and presence of saturating ligands. (**c**) FRET efficiencies of AF555/AF647 and Cy3/Cy7 pairs as a function of inter-dye distances calculated based on R0 values, 51 Å and 40.5 Å for AF555/AF647 and Cy3/Cy7 pairs, respectively. (**d**) smFRET population contour plots (top) and histograms (bottom) for Cy3/Cy7 labelled α_2A_AR -148C/367C in the absence and presence of saturating ligands. (**e**) smFRET distributions of AF555/AF647 labelled β2AR-148C/266C in apo, ISO-bound and ISO+Gs^GDP^ bound conditions. Black and purple lines represent Gaussians fitted to the high-FRET inactive and low-FRET active states, respectively. Dashed lines indicate the distinct mean FRET values. n represents the number of traces used to calculate the corresponding histograms. Data are mean ± s.e.m. from at least three groups of independent movies. Each group contains at least 3 movies. (**f**) Comparison of FRET peak centers for β2AR-148C/266C in different conditions. Data are mean ± s.e.m. from at least three groups of independent movies. (**g**) Example fluorescence (green for AF555; red for AF647) and FRET (blue) time traces for β2AR -148C/266C imaged in different conditions.

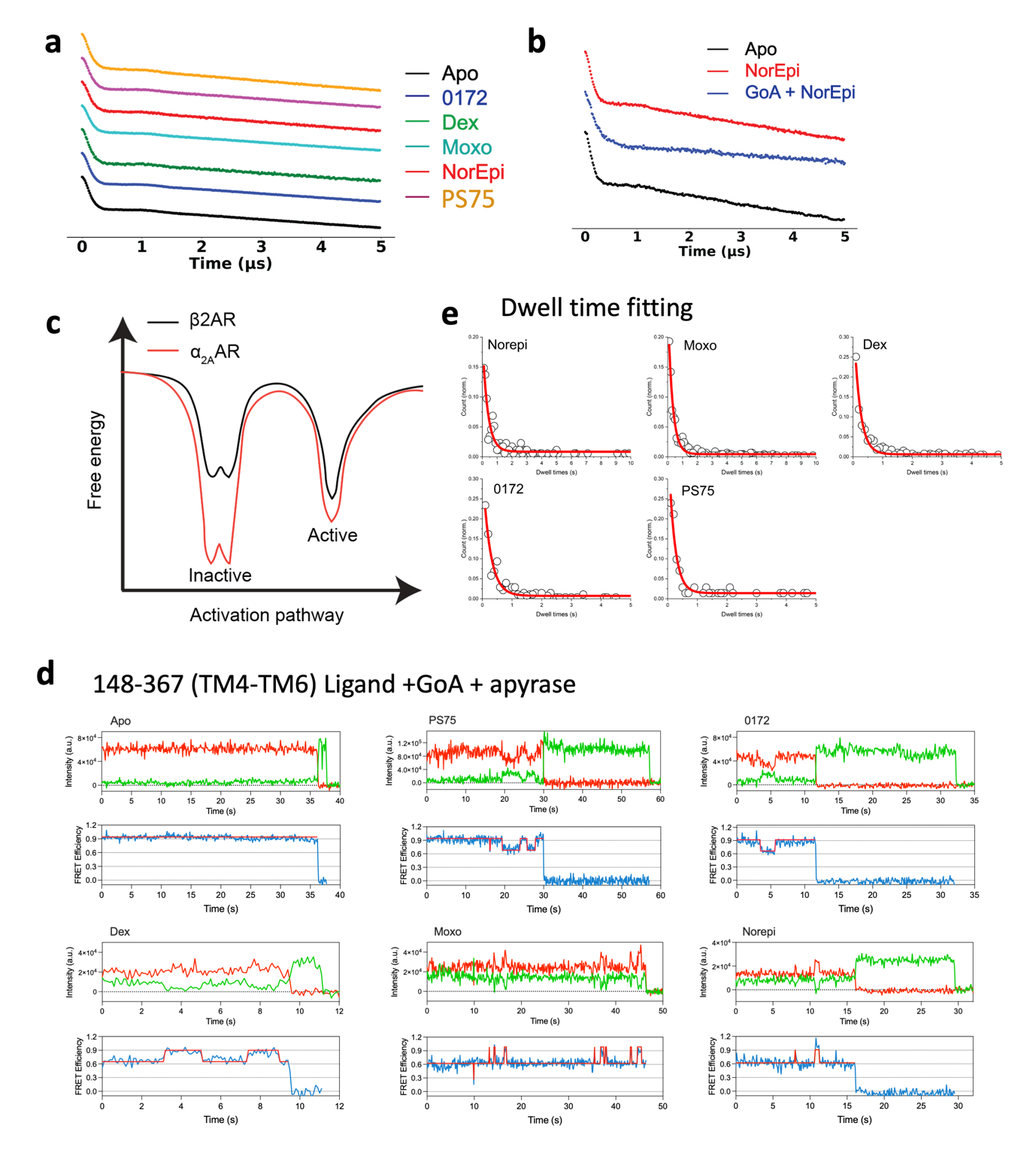
**Extended data Fig.3 TM6 conformational dynamics of α_2A_AR in the presence or absence of GoA.** (**a**) Dipolar evolutions for TM4-TM6 sensor (148/367) in apo and different ligand conditions. (**b**) Dipolar evolutions for TM4-TM6 sensor under different conditions: apo in gray, NorEpi in red and NorEpi+GoA^apyrase^ in light blue. (**c**) Comparison of energy landscapes for α_2A_AR and β2AR. (**d**) Example fluorescence (green for AF555; red for AF647) and FRET (blue) time traces for α_2A_AR -148C/367C imaged in different ligand conditions with GoA^apyrase^. Predicted state sequence was shown in red. (**e**) Fitting low-FRET dwell time. Cumulative counts are shown as black circles. low-FRET dwell times are fitted in single exponential decays (solid lines in red).

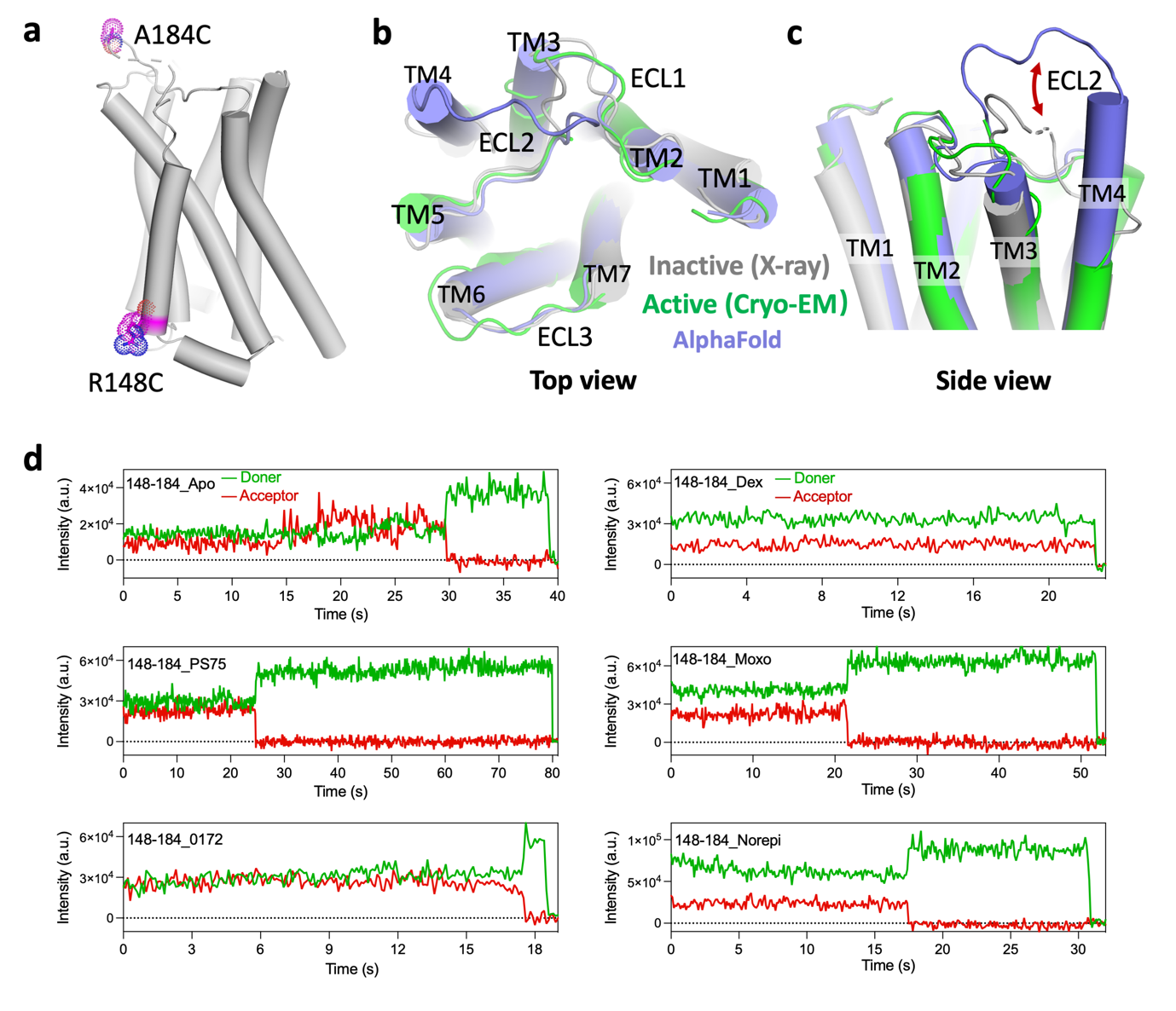

**Extended data Fig.4 Conformational dynamics of ECL2.** (**a**) Labeling sites of the TM4-ECL2 sensor. The cysteine mutations A184C and R148C were shown in dots. (**b-c**) Structural comparison of ECL2 in inactive and active states as well as in AlphaFold predicted model. (**d**) Example single-molecule fluorescence (green for AF555; red for AF647) traces of TM4-ECL2 sensor are shown for each ligand condition. The corresponding FRET traces were shown in Fig. 3c.

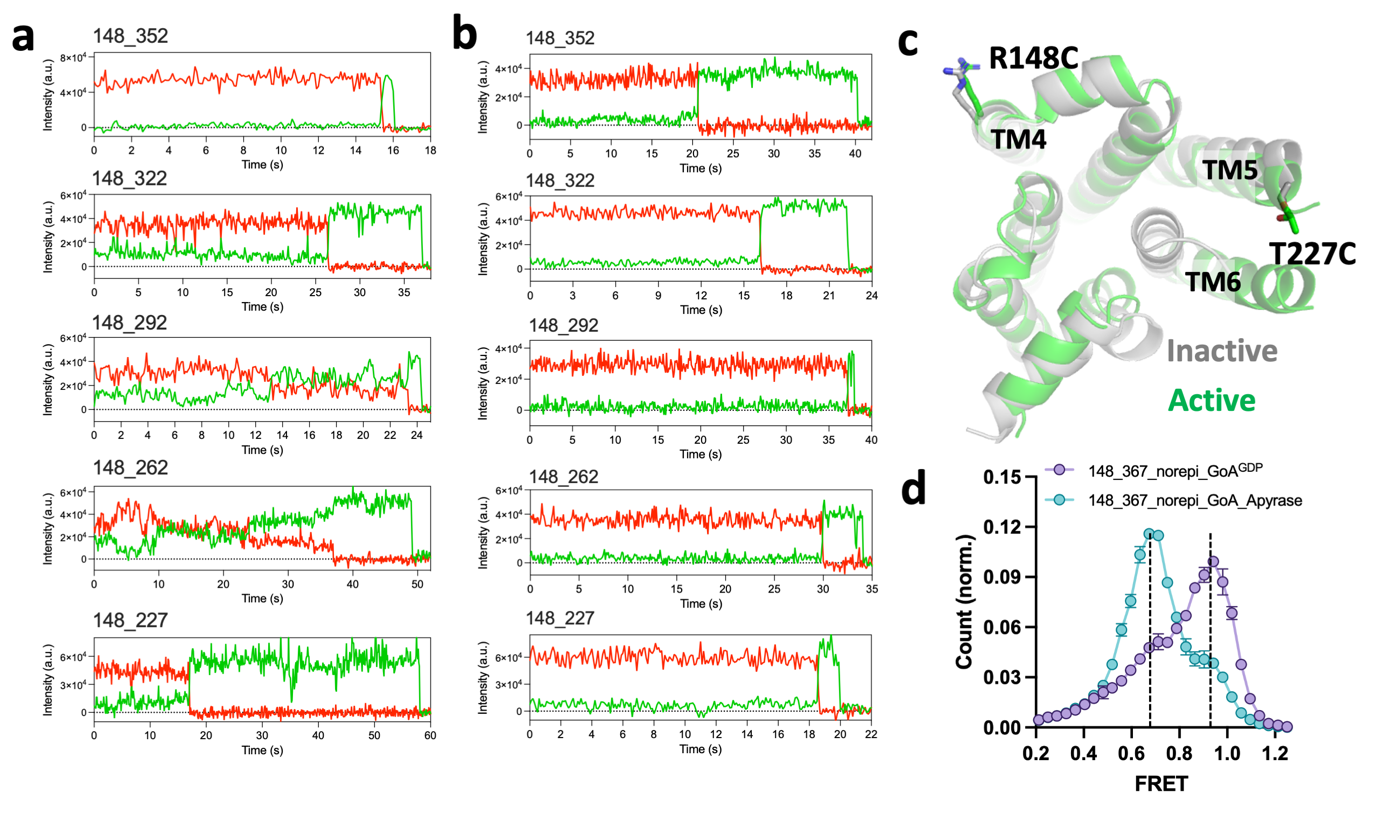

**Extended data Fig.5 Conformational dynamics of ICL3.** (**a-b**) Two example single-molecule fluorescence (green for AF555; red for AF647) time traces for TM4-ICL3 sensors in Apo state. The corresponding FRET traces were shown in **Fig. 4d-e**. (**c**) Structural comparison of T227C in inactive and active states. (**d**) Overlaid smFRET histograms of the α_2A_AR TM4-TM6 sensor (148/367) in the presence of norepi+GoA^GDP^ and norepi+GoA^apyrase^. Data are mean ± s.e.m. from at least three groups of independent movies. Each group contains at least 3 movies.

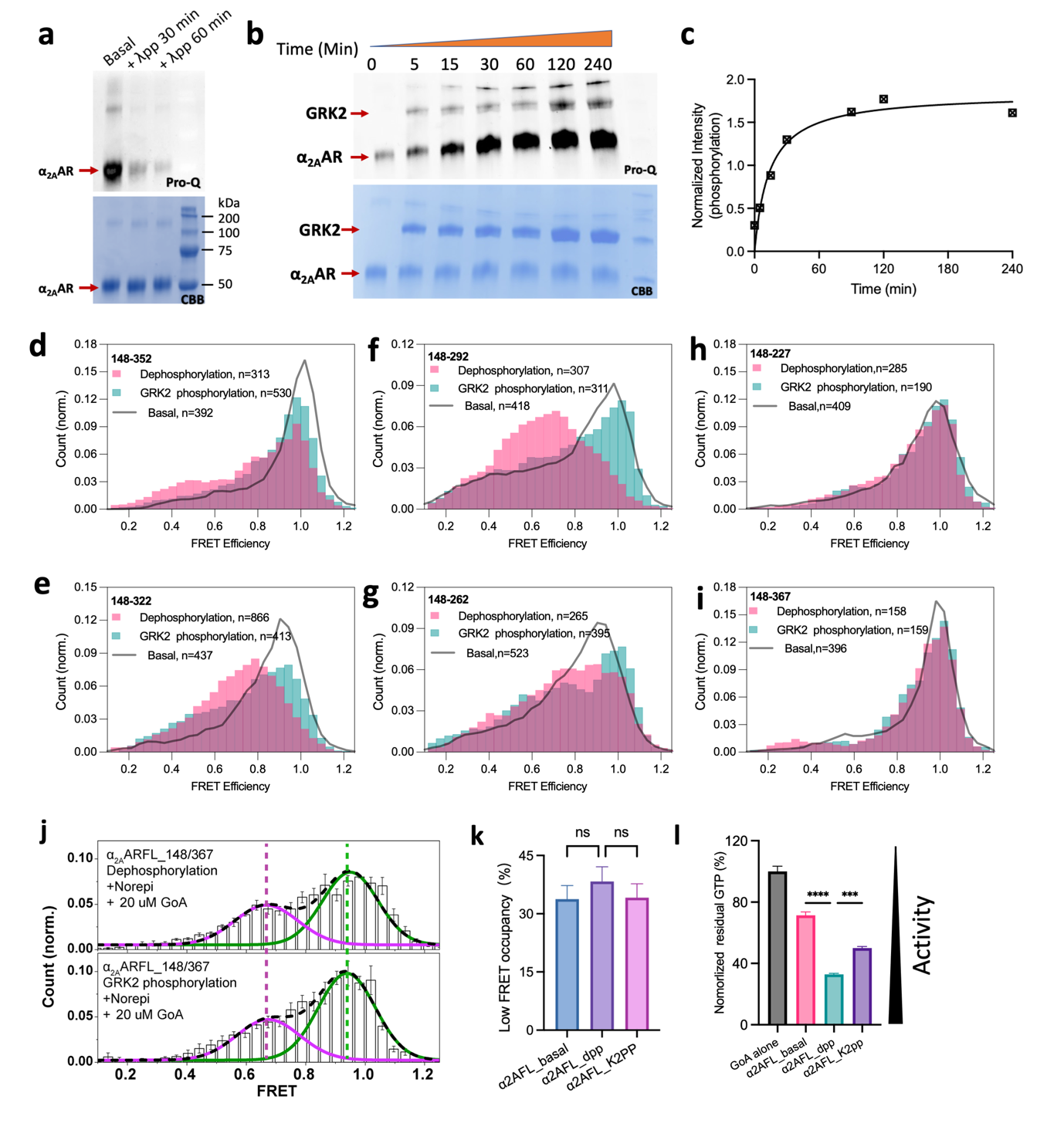
**Extended data Fig. 6 Effect of phosphorylation on ICL3 conformational dynamics. (a-b)** SDS-PAGE for dephosphorylation of the α_2A_AR by lambda phosphatase(λpp) (**a**) and phosphorylation by GRK2 (B) at different times. The same gel was stained by Pro-Q (upper) and Coomassie blue (bottom) sequentially. **(c)** Normalized intensities of time-dependent GRK2 phosphorylation extracted from (**b**). **(d-i)** Overlaid smFRET distributions of TM4-ICL3 sensors in the dephosphorylated (pink) and GRK2 phosphorylated (green) states. The distributions of basal-phosphorylated receptors were shown in gray as a reference. n represents the number of traces used to calculate the corresponding histograms. (**j**) smFRET distributions of the TM4-TM6 sensor (148/367) in the presence of Norepi and 20 uM GoA^GDP^ for dephosphorylated (upper) and GRK2 phosphorylated (bottom) receptors. Green and purple lines represent Gaussians fitted to the high-FRET inactive and low-FRET active states, respectively. Green and purple dash lines indicate the distinct mean FRET values. Black dash line represents the cumulative fitted distributions. Data are mean ± s.e.m. from at least three groups of independent movies. Each group contains at least 3 movies. (**k**) Low-FRET state occupancies calculated from panel (**j**). (**l**) GTP turnover activity for the basal-phosphorylated, dephosphorylated and GRK2 phosphorylated receptors. Errors represent mean ± s.e.m.

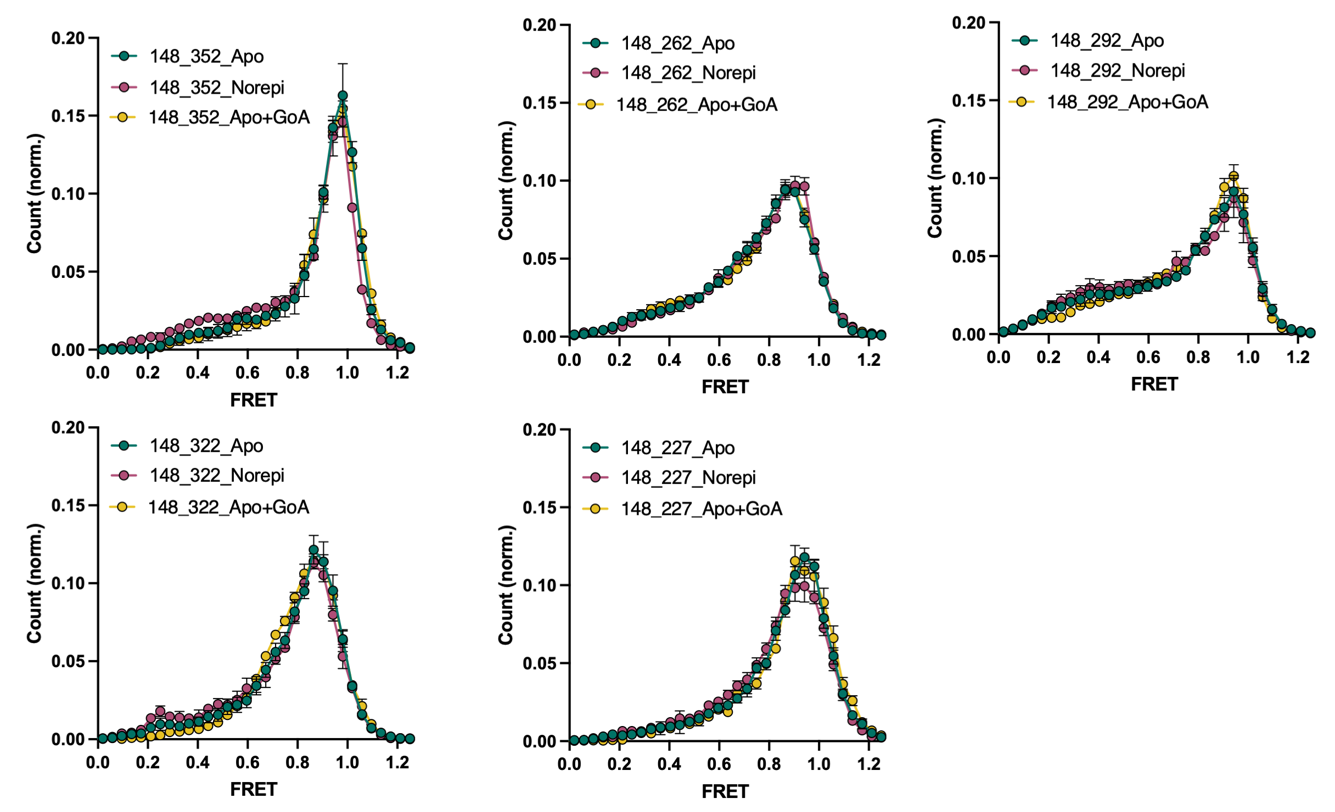

**Extended data Fig. 7 smFRET histograms of different ICL3 sendors.** Overlaid smFRET histograms for five TM4-ICL3 sensors in the Apo, Norepi-bound and Apo+GoA^apyrase^ conditions. Data are mean ± s.e.m. from at least three groups of independent movies. Each group contains at least 3 movies.

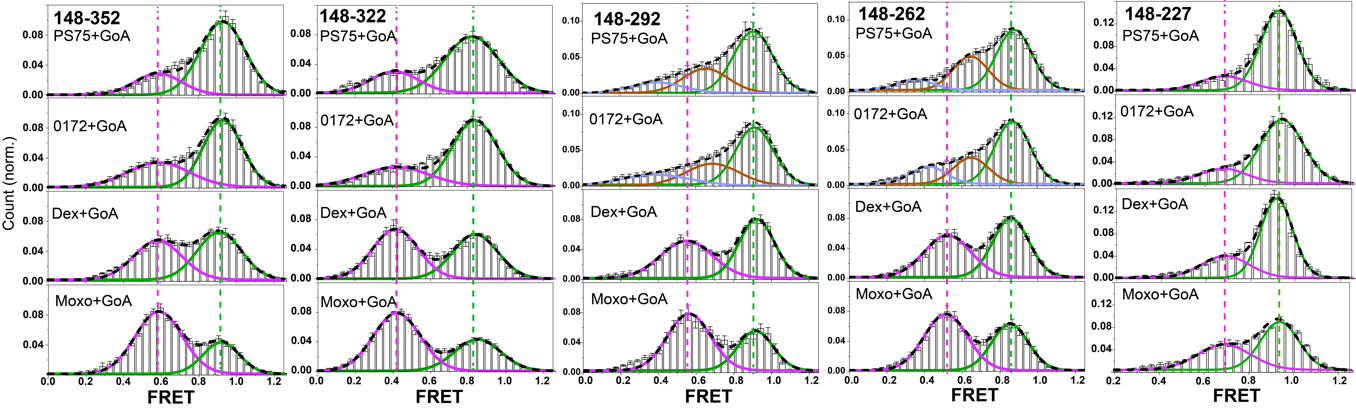

**Extended data Fig. 8 Ligand-specific conformatonal landscape of ICL3.** smFRET distributions of the TM4-ICL3 sensors in the presence of difference agonists + GoA treated with apyrase. Green and purple lines represent Gaussians fitted to the high-FRET inactive and low-FRET active states, respectively. Green and purple dash lines indicate the distinct mean FRET values. Black dash line represents the cumulative fitted distributions. Data are mean ± s.e.m. from at least three groups of independent movies. Each group contains at least 3 movies.

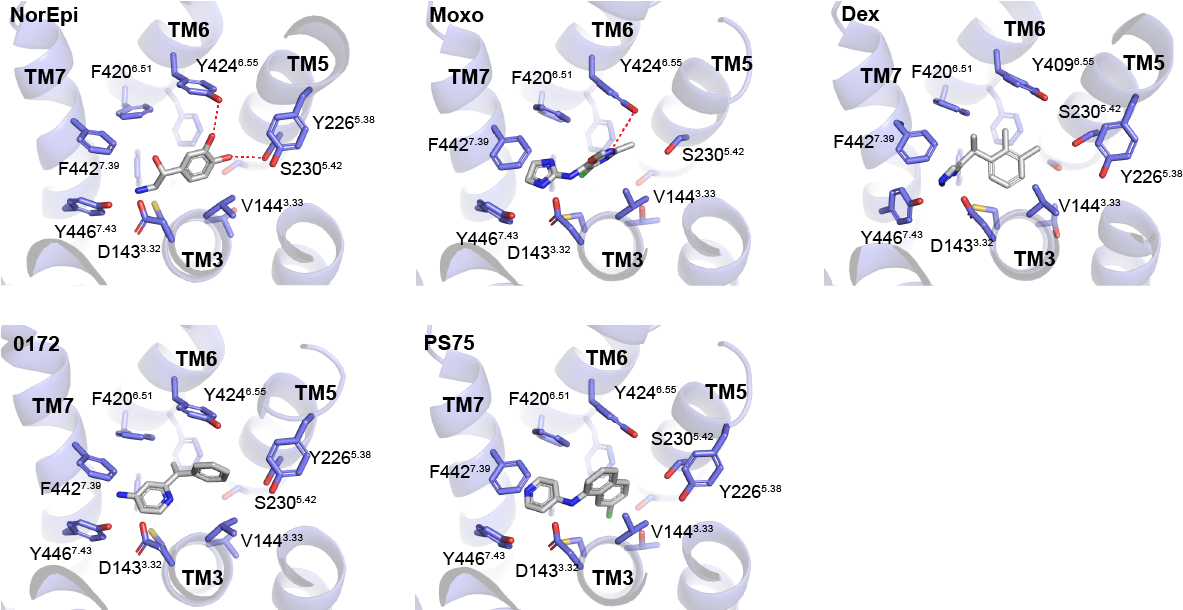

**Extended data Fig. 9 Agonist binding mode.** Comparison of agonist binding modes from cryo-EM structure (NorEpi, PDB:7EJ0; and Dex PDB:7EJA) or molecular docking (Moxo, 0172 and PS75).
